# A natural single nucleotide mutation in the small regulatory RNA ArcZ of *Dickeya solani* switches off the antimicrobial activities against yeast and bacteria

**DOI:** 10.1101/2021.07.19.452942

**Authors:** T Brual, G Effantin, J Baltenneck, L Attaiech, C Grosbois, M Royer, J Cigna, D Faure, N Hugouvieux-Cotte-Pattat, E Gueguen

**Author notes:** These authors contributed equally to this work (T Brual, G Effantin).

## Abstract

The necrotrophic plant pathogenic bacterium *Dickeya solani* emerged in the potato agrosystem in Europe. All isolated strains of *D. solani* contain several large polyketide synthase/non-ribosomal peptide synthetase gene clusters. Analogy with genes described in other bacteria suggests that the clusters *ooc* and *zms* are involved in the production of secondary metabolites of the oocydin and zeamine families, respectively. A third cluster that we named *ssm* for *<u>s</u>olani* <u>s</u>econdary <u>m</u>etabolite had an unknown function. In this study, we constructed mutants impaired in each of the three secondary metabolite clusters *ssm, ooc*, and *zms* to compare first the phenotype of the *D. solani* wild-type strain D s0432-1 with its associated mutants. We demonstrated the antimicrobial functions of these three PKS/NRPS clusters against bacteria, yeasts or fungi. The secondary metabolite cluster *ssm*, conserved in several other *Dickeya* species, produces a secondary metabolite inhibiting yeasts. Phenotyping and comparative genomics of different *D. solani* wild-type isolates revealed that the small regulatory RNA ArcZ plays a major role in the control of the clusters *ssm* and *zms*. A single-point mutation, conserved in some *Dickeya* wild-type strains, including the type strain IPO 2222, impairs the ArcZ function by affecting its processing into an active form.

**AUTHOR SUMMARY:** The development of new antibacterial molecules is critical in tackling the emergence of new pathogens or bacterial strains resistant to already available antibiotics. Bacterial phytopathogens can potentially synthesize novel compounds capable of targeting a specific type of microorganism. *Dickeya solani* is the only of the twelve described *Dickeya* species that has the three secondary metabolic pathways *ssm, ooc* and *zms*. An investigation of the functions of these three clusters allowed us to identify the anti-yeast activity of the *ssm* cluster, a potential new molecule of clinical importance. By comparing the antimicrobial activity of several *Dickeya solani* strains, we identified the small RNA regulator ArcZ as a critical regulator in the activation of the *ssm* and *zms* clusters. Our study showed that single-nucleotide polymorphisms of sRNA encoding genes can have huge impacts on bacterial phenotypes. It is thus critical to pay attention to the allele diversity of sRNA genes.

## INTRODUCTION

Bacterial phytopathogens of the genus *Dickeya* and *Pectobacterium* are pectinolytic necrotrophic bacteria with a broad host plant spectrum [1,2]. These members of the family *Pectobacteriaceae* [3] cause substantial agricultural losses worldwide by affecting many vegetables, ornamentals and crops, of which the potato is the most important economically. These bacteria are able to invade and degrade the plant tissues through the coordinated expression of genes encoding virulence factors, with a major role of pectate lyases that dissociate the plant cell wall constituents [4].

The *Dickeya* genus was established in 2005 [5], resulting from the reclassification of *Pectobacterium chrysanthemi* (formerly *Erwinia chrysanthemi*). To date, twelve species of *Dickeya* have been described, *D. aquatica, D. chrysanthemi, D. dadantii, D. dianthicola, D. fangzhongdai, D. lacustris, D. oryzae, D. parazeae, D. poaceiphila, D. solani, D. undicola*, and *D. zeae* [5–14].

The species *D. solani* was officially established in 2014 [9] but *D. solani* isolates have attracted attention since its emergence on the potato agrosystem in Europe in the early 2000s. It causes symptoms in both subtropical and temperate climates. Many scientific efforts have been made to provide information on this phytopathogen, resulting in 76 *D. solani* genomes available in May 2021 [15]. Comparative genomics was performed to identify the genetic basis for the different levels of virulence between *D. solani* strains [15–20]. Most *D. solani* strains isolated from different regions show a low level of genetic variation, suggesting a clonal origin [20]. The *D. solani* genomes share a high similarity and synteny with those of the model strain *D. dadantii* 3937, prompting comparison between the two species. Only a few hundred genes were specific to each species, including a few dozen distinctive genomic regions [16,17]. Three of these regions encode polyketide synthases (PKS), non-ribosomal peptide synthetases (NRPS) and amino acid adenylation domain proteins, which are typically involved in the production of secondary metabolites [16].

PKSs and NRPSs are able to synthesize molecules by sequential condensation of carboxylic acids and amino acids, respectively. PKS and NRPS modules can combine together to form hybrid PKS/NRPS systems capable of producing compounds of great structural diversity [21]. The molecules synthesized may have siderophore, antibiotic or phytotoxic properties that promote the virulence of a plant pathogen. Three PKS/NRPS clusters are present in all sequenced *D. solani* strains and found in a few other *Dickeya* species and related genera: *ooc, zms*, and a cluster of unknown function [22]. In *Serratia plymuthica*, the cluster *ooc* is involved in the synthesis of oocydin A, a halogenated macrolide with antifungal and anti-oomycete activity [23]. The cluster *zms*, previously found in the genomes of *S. plymuthica* and *Dickeya oryzae*, leads to the biosynthesis of a polyamino-amide antibiotic, zeamine [24,25]. The third cluster *ssm* (for *<u>s</u>olani* <u>s</u>econdary <u>m</u>etabolite) is found in a few *Dickeya* species but the nature and function of the synthesized molecule are unknown. To elucidate the contribution of the *ssm, ooc* and *zms* clusters of *D. solani* in competition with other living organisms, we constructed mutants of each of the three loci involved in the biosynthesis of secondary metabolites in the highly virulent *D. solani* strain D s0432-1. We showed that the *D. solani* D s0432-1 clusters *zms* and *ooc*, encoding zeamine and oocydin biosynthesis, are involved in growth inhibition of bacteria and fungi, respectively. Recently we, and others identified the third cluster as implicated in *Dickeya solani*’s ability to inhibit the growth of different yeast species [26,27]. We named this cluster *ssm* (for *solani* secondary metabolite) while Matilla and colleagues named it sol (for solanimycin, the molecule produced by this cluster). Finally, the comparative analysis of the genomes and phenotypes of several wild-type *D. solani* strains revealed the fundamental role of the small RNA ArcZ. This sRNA plays a major role in the regulation of the *ssm* and *zms* clusters, which can be disrupted by a mutation in the *arcZ* sequence, conserved in some natural isolates.

## RESULTS

### Description and distribution of the three selected PKS/NRPS secondary metabolite clusters of *D. solani*

The gene clusters encoding complex NRPS and PKS involved in the production of secondary metabolites were named *ssm, ooc* and *zms*, respectively (Fig. 1A). Their distribution in *Dickeya* species is summarized in figure 1B (see also Table S1 for a complete list of the *Dickeya* strains with the indication of the presence or absence of each cluster).

**Figure 1.**
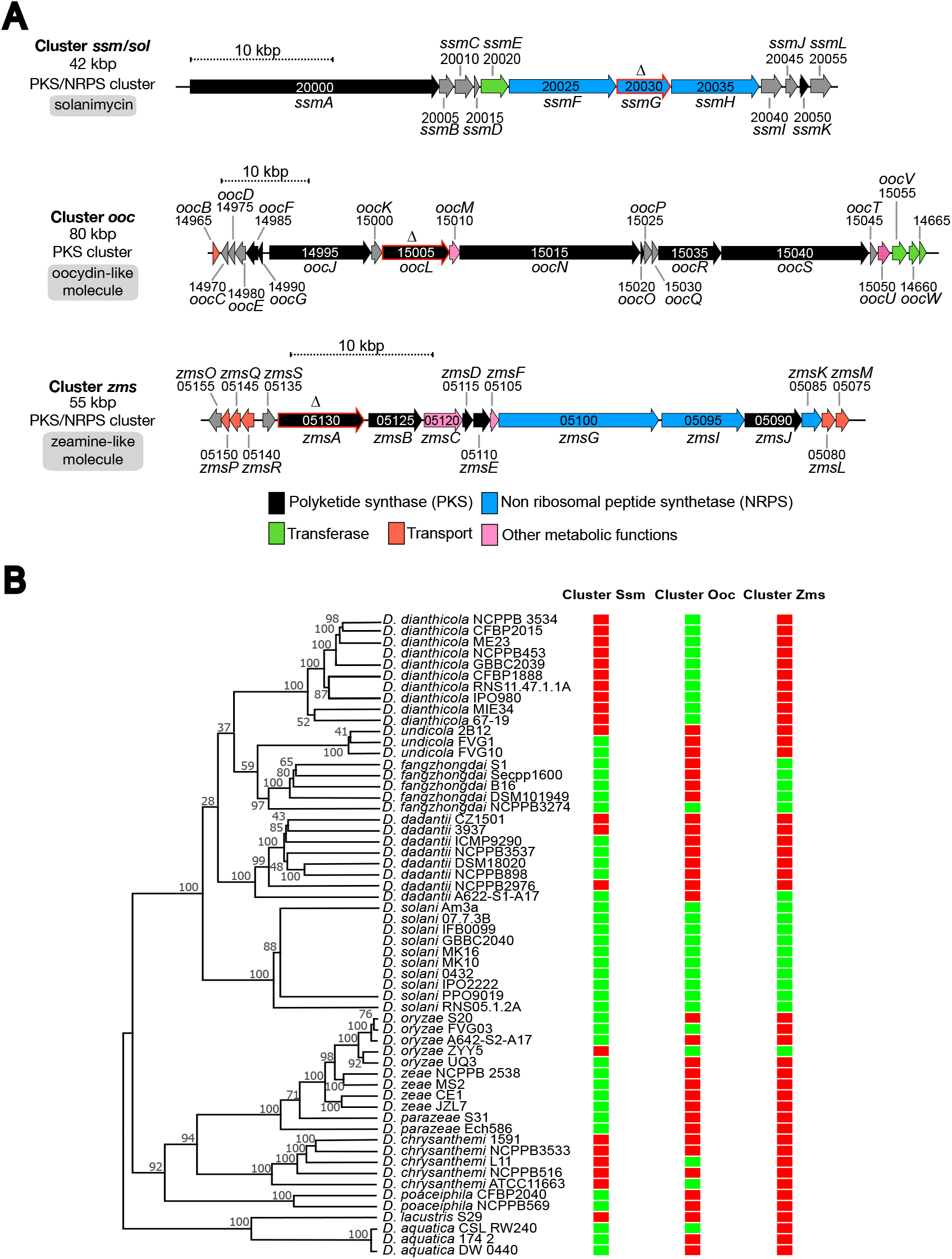
Organization and distribution of the secondary metabolite clusters *ssm, ooc*, and *zms* of *D. solani* D s0432-1. (A) Organization of the clusters *ssm, ooc*, and *zms*. Genes are indicated using the NCBI nomenclature of the NCBI reference genome sequence NZ_CP017453.1 of *D. solani* D s0432-1. XXXXX are the digital number in BJD21_RSXXXXX, which corresponds to the locus tag. Arrowheads show gene orientations. Color code indicates gene function. The red-framed arrows indicate genes targeted for in-frame deletion performed in this study. (B) MLSA tree positioning strains within the *Dickeya* genus. The evolutionary history was inferred by using the Maximum Likelihood method. The presence (green frame) or absence (red frame) of the cluster *ssm, ooc*, and *zms* are indicated in front of each strain. The distribution of these clusters in 155 *Dickeya* strains whose genome has been sequenced is given in Table S1.

The ~42-kbp cluster *ssm* contains the 12 genes *ssmABCDEFGHIJKL* (Fig. 1A). It is widely conserved in the genus *Dickeya, i*.*e*., in all sequenced *D. solani, D. aquatica, D. fangzhongdai*,

*D. poaceiphila* and *D. zeae* genomes, in some *D. dadantii* strains (such as NCPPB 898, NCPPB 3537, DSM18020, but not the model strain 3937), in some *D. undicola* strains (FVG1, FVG10), and in some *D. oryzae* strains (such as S20 and FVG03). The structure of the metabolite produced from this cluster has not been elucidated yet but the function of this cluster has been described by us in a 2021 preprint article [27] and others in 2022 [26]. The *ssm* cluster produces solanimycin that targets yeast.

The ~80-kbp cluster *ooc* (Fig. 1A) is highly similar to the *oocBCDEFGJKLMNOPQRSTUVW* cluster of *S. plymuthica* A153. It is present in all the sequenced genomes of *D. solani* and *D. dianthicola*, in some *D. oryzae* strains (such as ZYY5, EC1, DZ2Q and ZJU1202) and in a few other species. In *S. plymuthica* A153, disruption of this gene cluster abolished bioactivity against the fungi *Verticillium dahliae* and the oomycetes *Pythium ultimum*. This cluster produces oocydin A [23], a chlorinated macrolide, powerfully active against plant pathogenic oomycetes [28]. Since various *D. solani* strains inhibit *V. dahliae* and *P. ultimum* growth [29], it was suggested, on the basis of gene sequence homologies and similar cluster organization, that *D. solani* also produces oocydin A.

The ~55-kbp cluster *zms* encodes mixed fatty acid synthase FAS/PKS and hybrid NRPS/PKS enzymes (Fig.1A). Its genomic organization is identical to the *D. oryzae* EC1 zeamine cluster and related to the *S. plymuthica* AS12 zeamine cluster [24,30]. While the *zmsABCDEFGIJKLMNPQRS* cluster directing zeamine biosynthesis is present only in a few *D. oryzae* strains, it is conserved in all the sequenced *D. solani* and *D. fangzhongdai* genomes, which suggest secondary acquisition by horizontal gene transfer in *D. oryzae* [22,30]. After reclassification of several *D. zeae* strains in the novel species *D. oryzae* [14], the *zms* cluster appeared to be absent in the genomes of true *D. zeae* strains. It is present in some, but not all, *D. oryzae* rice strains (ZYY5, EC1, DZ2Q and ZJU1202) and in only one *D. dadantii* strain (A622-S1-A17). The zeamine biosynthetic clusters from *D. oryzae* EC1 and *D. solani* Ds0432-1 share from 59 to 94% identity at individual protein level [30]. Zeamine-related antibiotics are polyamino-amide molecules toxic to a wide range of pro- and eukaryotic organisms such as bacteria, fungi, oomycetes, plants, and nematodes [31]. Mutation of the zeamine synthase gene *zmsA* in *D. oryzae* EC1 attenuates the inhibition of rice seed germination [24] and suppresses antibacterial activity against *E. coli* [24]. Zeamine produced by *S. plymuthica* kills nematodes and yeast [32]. *D. solani* IPO 2222 can also kill the nematode *Caenorhabditis elegans* but not as quickly as *S. plymuthica* [32].

To interrupt the synthesis of the secondary molecules produced by these three clusters, we constructed from a Nal^R^ Gm^R^ derivative of WT *D. solani* D s0432-1 in-frame deletion mutants inactivating a key gene of each cluster. The inhibitory effects of the mutants Δ*ssmG*, Δ*oocL* and Δ*zmsA* against fungi, bacteria and yeasts were compared with that of the parental strain. To further analyze possible synergistic effects, double mutants Δ*ssmG*Δ*oocL*, Δ*ssmG*Δ*zmsA* and Δ*oocL*Δ*zmsA* as well as the triple mutant (hereafter called Δ3) were also constructed.

### The *D. solani* oocydin cluster ooc inhibits *Ascomycota* growth, and the *ssm* cluster is also involved in the inhibition of some fungi

We compared the ability of the WT *D. solani* strain D s0432-1 and its derived mutants to inhibit the growth of *Botrytis cinerea, Magnaporthe oryzae* and *Sclerotinia sclerotiorum*, three fungi-like eukaryotes of the phylum *Ascomycota*. The center of a potato dextrose agar (PDA) plate was inoculated with fungal mycelium, and 5 µl of overnight bacterial culture of each *D. solani* strain grown in M63 minimal medium with sucrose was deposited at the periphery of the plate (Fig. 2). The different *D. solani* strains showed similar overall growth in liquid culture (Fig. S1). After incubation at 25°C for several days, we observed a growth inhibition of the three fungi by the *D. solani* WT and the Δ*ssmG* and Δ*zmsA* mutants. In contrast, the Δ*oocL* mutant seemed less effective: *B. cinerea* and *M. oryzae* mycelium growth was completely unaffected, while a small zone of inhibition was observed for *S. sclerotiorum* (Fig. 2). This suggests that the Δ*oocL* mutant may also produce another molecule that inhibits this fungus. The phenotypes were fully restored in the reverted strain Δ*oocL/oocL*^*+*^. Double and triple mutants of the secondary metabolite clusters containing the *oocL* deletion showed no differences from the Δ*oocL* mutant with *B. cinerea* and *M. oryzae*. However, the Δ*oocL*Δ*ssmG* double mutant and the Δ3 triple mutant were completely unable to inhibit *S. sclerotiorum* growth (Fig. 2). These data suggest that the *ooc* cluster is mainly responsible for the majority of *Ascomycota* inhibition and that the *ssm* cluster is secondary involved in the inhibition of some fungi. In contrast, the *zms* cluster does not seem to be involved in fungal inhibition.

**Figure 2.**
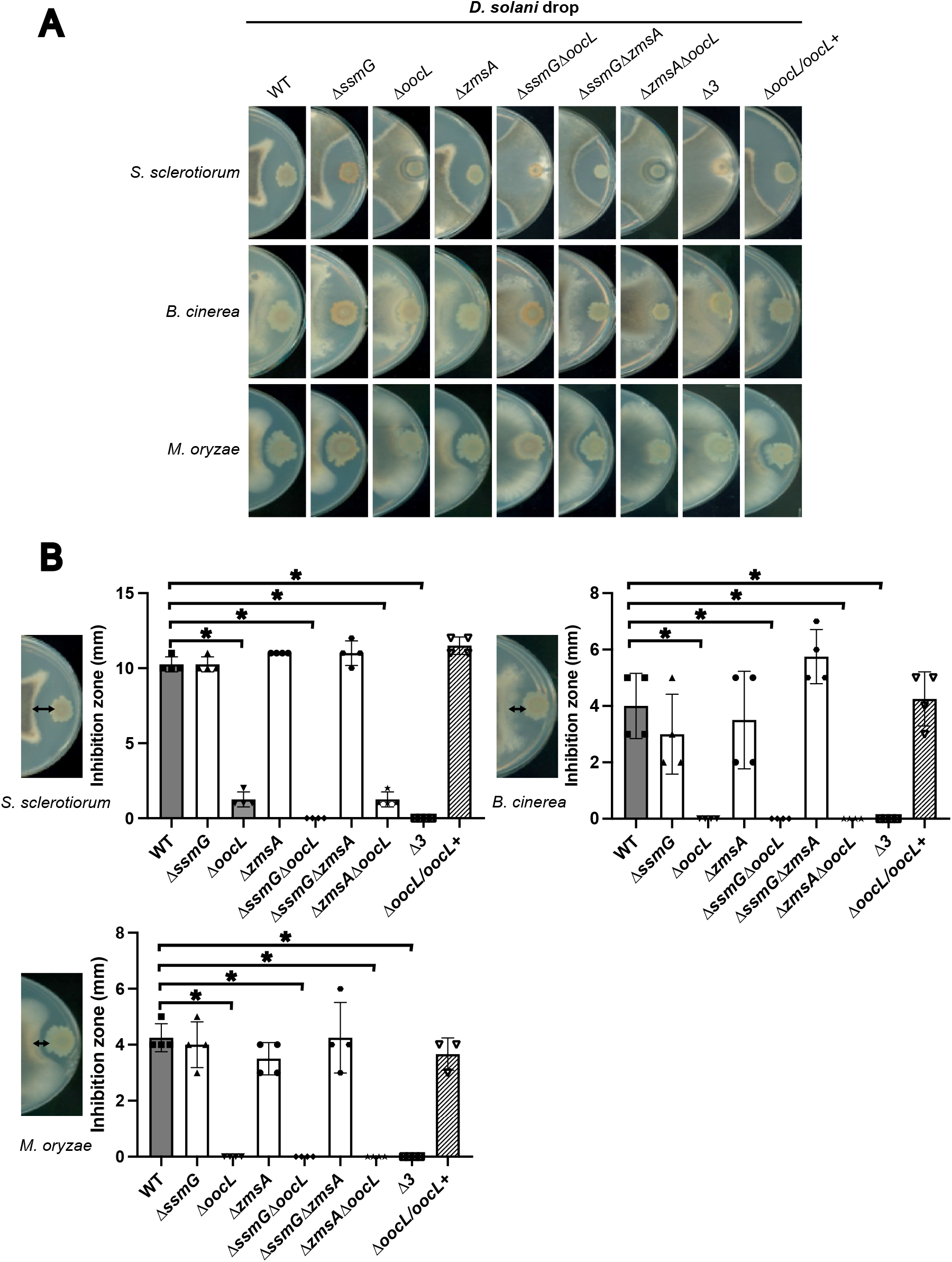
Inhibition of fungal growth by *D. solani* Ds0432-1 and mutant derivatives. 5 µL of bacteria culture at OD_600nm_= 2 from *D. solani* Ds0432-1 (WT) or derivatives were spotted onto PDA plates inoculated with plugs of *S. sclerotiorum, B. cinerea or M. oryzae mycelium*. Plates were incubated at 25°C until the mycelium covers the plate. Lengths of fungi inhibition zone were measured in 4 independent experiments. A statistical difference was significant only between the WT and Δ*oocL* mutants (Mann-Whitney test; * p-value < 0.05).

### The *D. solani* zeamine cluster inhibits bacterial growth

The zeamine cluster of *D. oryzae* EC1 is responsible for the a bactericidal activity against *Escherichia coli* DH5*α* [24]. *D. oryzae* EC1 and *D. solani* zeamine biosynthetic genes share a high degree of similarity [30]. We thus evaluated the capacity of the *D. solani* strain D s0432-1 and its derived mutants to inhibit the growth of gram-positive and gram-negative bacteria (Fig. 3). Only the Δ*zmsA* mutant was unable to inhibit *Bacillus subtilis* growth, indicating that WT *D. solani* produces an active zeamine antibiotic (Fig. 3). The WT phenotype was restored in the reverted Δ*zmsA*/*zmsA+* strain. We also tested the ability of *D. solani* to inhibit growth of the other gram-positive bacteria *Streptomyces scabiei*, a plant pathogen causing the potato disease common scab [33]. An inhibition of *S. scabiei* growth was observed, except with the Δ*zmsA* mutant. No phenotype difference was observed between the single Δ*zmsA* mutant and the double and triple mutants containing the *zmsA* deletion, suggesting that only the cluster *zms* is required for Gram-positive bacterial growth inhibition. We also tested the ability of *D. solani* to inhibit growth of the gram-negative bacteria *E. coli, D. dadantii* and *Pectobacterium atrosepticum* (Fig. 3). *E. coli* DH5*α* was, like *B. subtilis* and *S. scabiei*, inhibited by the WT strain through *zms* activity, albeit much less potently (Fig. 3 and Fig. S2). No inhibition of the two other pectinolytic bacteria *D. dadantii* and *P. atrosepticum* growth was observed. In conclusion, out of the three clusters analyzed, only the zeamine-like molecule potentially produced by the *zms* cluster has an anti-bacterial activity against some gram-positive and gram-negative bacteria.

**Figure 3.**
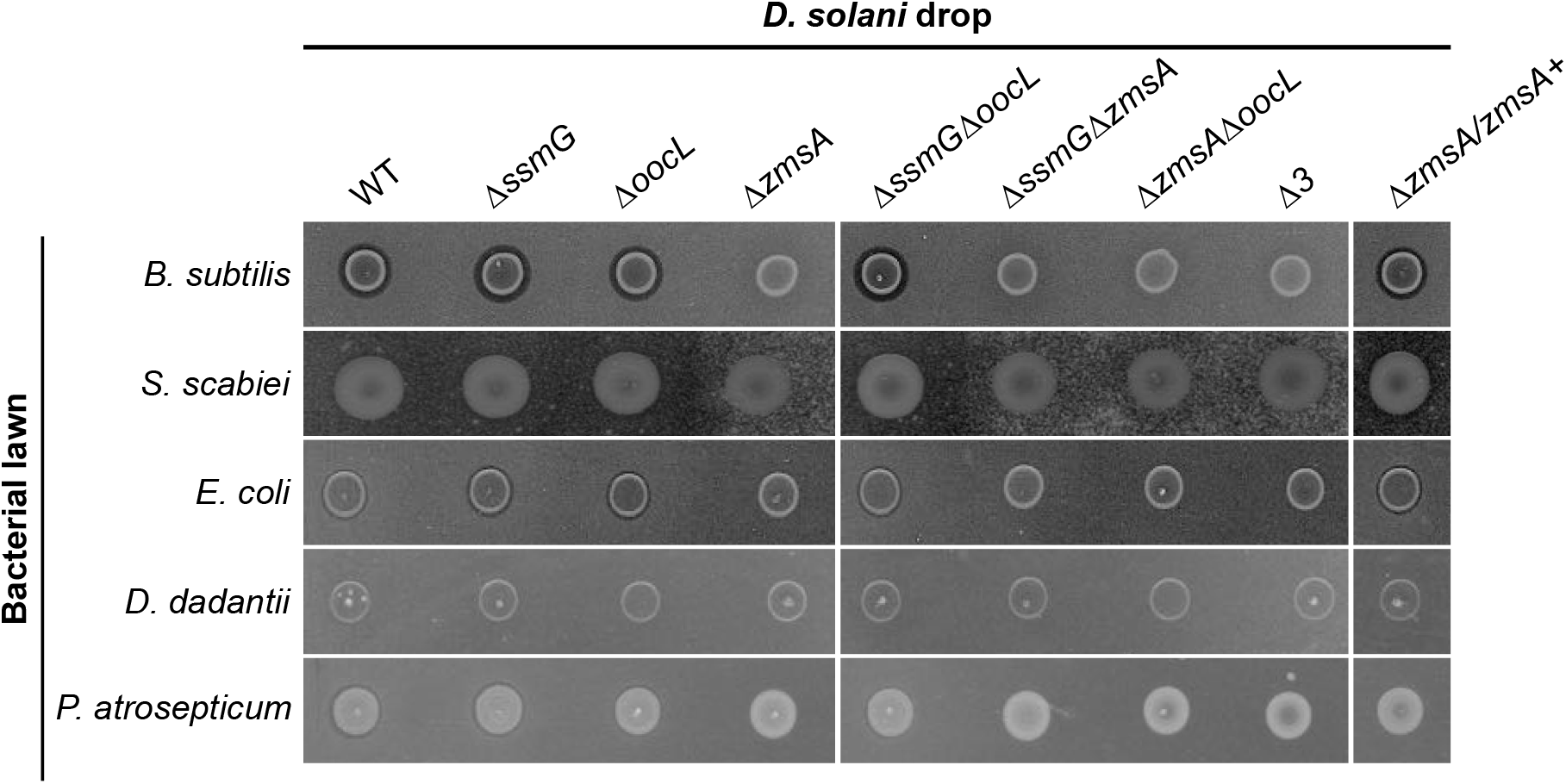
Inhibition of bacteria by *D. solani* Ds0432-1 and mutant derivatives. Bioassay plates were prepared by mixing bacterial culture (*B. subtilis, S. scabiei, E. coli, D. dadantii and P. atrosepticum*) with warm LB agar. 5 µL of bacterial cultures at OD_600nm_= 2 from *D. solani* Ds0432-1 (WT) or derivatives were spotted onto the plate and incubation was performed at 30°C during 48 h. A slight inhibition zone was observed with *B. subtilis* and *E. coli*, except with the *Δzms* mutants. All experiments were carried in 4 replicates.

### The *D. solani* clusters *ssm* and *zms* inhibits yeast growth

Since zeamine produced by *S. plymuthica* A153 has previously been shown to be toxic to the ascomycete yeast *Saccharomyces cerevisiae* [32], we tested the capacity of *D. solani* D s0432-1 and its derivative mutant to inhibit the growth of the yeasts *S. cerevisiae, Kluyveromyces lactis*, a predominant eukaryote during cheese productions [34], and *Candida albicans*, an opportunistic human pathogen. In a growth inhibition assay performed in Yeast Peptone Dextrose (YPD) solid agar medium, we observed a strong inhibition of *K. lactis* by WT *D. solani*, a lower inhibition of *S. cerevisiae*, and a slight inhibition of *C. albicans* (Fig. 4). The Δ*ssmG* mutant was almost completely deficient in its ability to inhibit yeast growth, while the Δ*ssmG*/*ssmG*^*+*^ revertant strain showed a restored WT phenotype. The cluster *ssm* is thus primarily responsible for the anti-eukaryotic action against these yeasts (Fig. 4). Nevertheless, a weak halo of inhibition of the yeasts *K. lactis* and *S. cerevisiae* was observable around the mutant Δ*ssmG*. This halo disappeared around the double mutant Δ*ssmG*Δ*zmsA* and the triple mutant *Δ3*, suggesting that the *zms* cluster is also implicated in yeast inhibition, but at a lower level than that observed for the *ssm* cluster. In conclusion, although the zeamine cluster of *D. solani* D s0432-1 is involved in the inhibition of yeast growth, as shown for *S. plymuthica* A153 [32], *D. solani* D s0432-1 possesses a second cluster *ssm* that produces another anti-yeast molecule.

**Figure 4.**
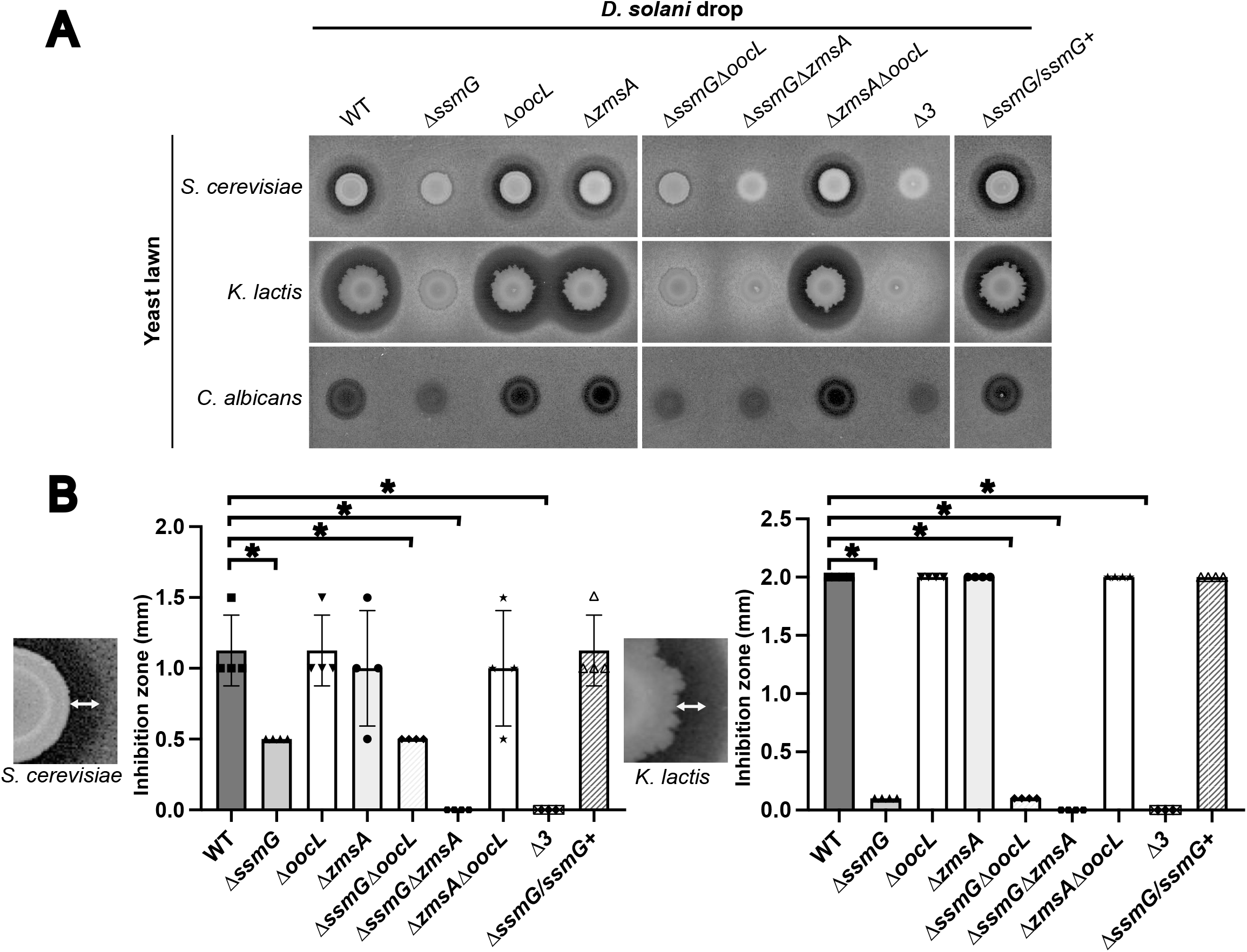
Inhibition of yeasts by *D. solani* Ds0432-1 and mutant derivatives. Bioassay plates were prepared by mixing yeast culture (*S. cerevisiae, K. lactis, C. albicans*) with warm YPD agar. (A) 5 µL of bacterial culture at OD_600nm_= 2 from *D. solani* Ds0432-1 (WT) or derivatives were spotted onto the plates and incubation was performed at 30°C during 48 h. (B) The radius of each inhibition zone was measured. A statistical difference was significant only between the WT and the *Δssm* mutants (Mann-Whitney test; * p-value < 0.05). For *C. albicans*, the inhibition zone was too small to be quantified. All experiments were carried in 4 replicates.

### The WT *D. solani* strains IPO 2222 and IFB0223 are unable to inhibit yeast and bacterial growth

We showed that *D. solani* D s0432-1 has the capacity to inhibit the growth of a wide range of microorganisms, including filamentous fungi, bacteria, and yeasts. We therefore tested whether other WT *D. solani* strains isolated from the environment in Europe, either from lesions of potato tubers or plants, or from potato rhizosphere, possess this ability. We focused our work on *D. solani* strains available in national strain collections and whose genome sequence is available. These strains are D s0432-1, IFB0099, IFB0158, IFB0223, IFB0484, IPO 3337, IPO 3494, IPO 3793 and the *D. solani* type strain IPO 2222. We tested their capacity to inhibit the bacterium *B. subtilis*, the yeast *K. lactis* and the fungus *S. sclerotiorum* (Fig. 5A). All strains were able to inhibit the growth of *S. sclerotiorum*. The inhibition of *B. subtilis* and *K. lactis* was clearly observed for all the strains except IPO 2222 and IFB0223. These strong phenotypic differences between IPO 2222 and IFB0223 and the other WT *D. solani* strains could not be explained by any nucleotide variabilities in the *ssm, zms*, and *ooc* clusters. All the selected strains carry the *ssm, zms*, and *ooc* clusters, with 100% conservation at the nucleotide level (including in the promoter regions). Nonetheless, there are certain changes at the genome level, most notably some nucleotide variability with the presence of SNPs (single-nucleotide polymorphism) and InDel (insertion or deletion), outside the clusters *ssm, zms*, and *ooc*. Hence, we searched for variations in the whole genomes by comparing two groups: strains IFB0223 and IPO 2222 that are defective for the growth inhibition of yeast and bacteria, and strains D s0432-1, IPO 3337 and IFB0099 that are efficient for these inhibitions (Table S2).

**Figure 5.**
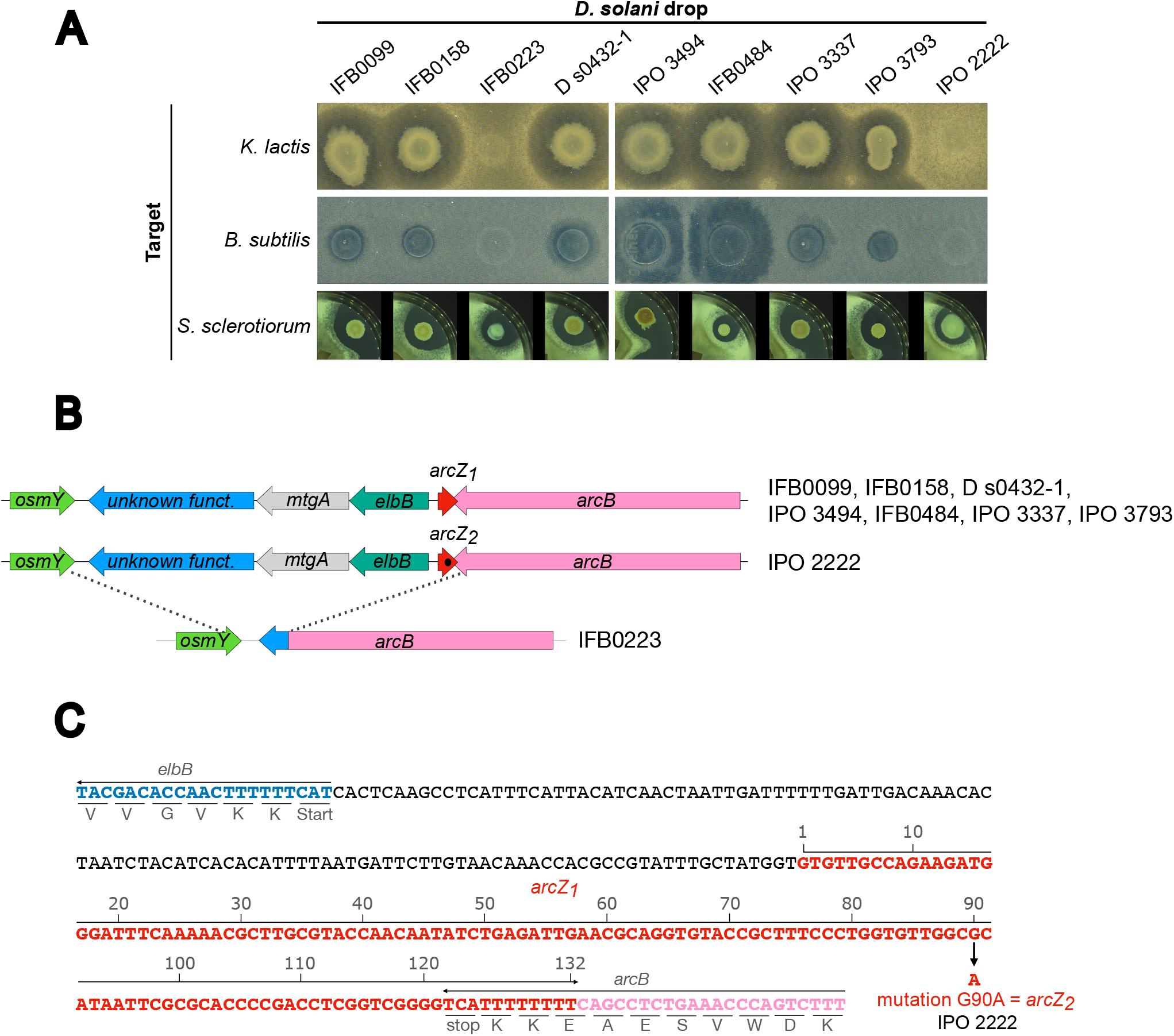
*D. solani* WT strains deficient for anti-microbial activity are mutated for the sRNA *arcZ*. (A) Inhibition of *B. subtilis, K. lactis* and S. *sclerotiorum* by diverse WT *D. solani* strains. The tests were performed as previously described using the nine strains IFB 0099 (NZ_CP024711), IFB 0158 (NZ_PENA00000000), IFB 0223 (NZ_CP024710), D s0432-1 (NZ_CP017453), IPO 3494 (NZ_CM001842), IFB 0484 (NZ_CM001860), IPO 3337 (NZ_CP016928), IPO 3793 (NZ_CP017454) or the type strain IPO 2222 (NZ_CP015137). Only IFB 0223 and IPO 2222 were unable to inhibit *B. subtilis* and *K. lactis*. (B) Organization and conservation of the *arcZ* locus in the nine *D. solani* strains. The single mutation G90A in IPO 2222 *arcZ* is symbolized by a dark spot in the red *arcZ* arrow. (C) Zoom on the nucleotide sequence around *arcZ* of *D. solani* D s0432-1. This allele is named *arcZ*_*1*_. Positions and orientations of the genes *elbB, arcZ* and *arcB* are indicated. The location of the G90A mutation present in IPO 2222 is indicated. This allele is named *arcZ*_*2*_.

A SNP was observed with an identical nucleotide in the strains D s0432-1, IPO3337 and IFB0099 (a G at position 90 of *arcZ*) but different nucleotide in the strain IPO2222 (a A at position 90 of *arcZ*) (position 2530087 according to IPO 2222 genome). This variation is positioned in the 3’ region of the small RNA *arcZ*, known to be a key area interacting with targeted mRNA for post-translational regulation in other bacterial species (Fig. 5C). The ArcZ variants of D s0432-1 and IPO 2222 will be referred to as ArcZ_1_ and ArcZ_2_ respectively in the remainder of this article. In the strain IFB0223, we observed a 3-kbp chromosomal deletion covering the gene *arcZ*, and extending between the genes *arcB* and *osmY* (Fig. 5B). In other *D. solani* strains, this region includes *arcZ, elbB, mtgA*, and a gene encoding an alginate lyase domain (pfam05426). These discoveries led us to hypothesize that the inability of strains IFB0223 and IPO 2222 to inhibit yeast and bacterial growth could be due to an ArcZ deficiency. Before going further, we ordered the *D. solani* IPO 2222 type strain stored in the BCCM collection (LMG25993) to analyse its genotype and phenotype. Using PCR amplification and sanger sequencing of *arcZ*, we confirmed that the *arcZ* allele of LMG 25993 is *arcZ*_2_, similarly to the IPO 2222 strain used in our experiments Similarly, the strain IPO 2222 used in our experiments and LMG 25993 have the same inhibition phenotype, i.e., an incapacity to inhibit yeast and bacterial growth (Fig. S3).

### The *arcZ* gene is required for *zms* and *ssm* anti-microbial activities

sRNAs are posttranscriptional regulators that most commonly influence gene translation by base-pairing to target mRNAs. In *E. coli*, this regulation often requires the RNA chaperones Hfq or ProQ [35–37]. ArcZ belongs to what is called the “core sRNAs” because it is highly conserved among enterobacterial species [38]. In order to determine if ArcZ is required for secondary metabolite production, the Δ*arcZ* mutants of *D. solani* strains D s0432-1 and IPO 2222 were constructed, and their phenotypes compared to those of the WT strains (Fig. 6A). In M63 sucrose, the mutants grew as well as the parental WT strains (Fig. S4). In our inhibition assay against *B. subtilis* and *K. lactis*, no inhibition zones were observed around the Δ*arcZ*_*1*_ mutant of D s0432-1 (Fig. 6A), demonstrating that ArcZ_1_ is required for the synthesis of the metabolites produced from the *ssm* and *zms* clusters in D s0432-1. However, the Δ*arcZ*_*1*_ mutant of *D. solani* D s0432-1 was still able to inhibit the growth of the fungus *S. sclerotiorum*, although a slight decrease in the anti-fungal effect was observed. Thus, ArcZ does not appear to be fully required for *ooc* cluster expression, which is consistent with the results obtained with *D. solani* IFB0223, the naturally *arcZ* deletion mutant capable of inhibiting *S. sclerotiorum* growth (Fig. 5A and Fig. 6A). *D. solani* IPO 2222 WT inhibited neither bacteria nor yeast but inhibited the growth of fungi, and the phenotype of the IPO 2222 Δ*arcZ*_*2*_ mutant was similar to that of the IPO 2222 WT or the D s0432-1 Δ*arcZ*_*1*_ mutant. This suggests that ArcZ_2_ might be inactive.

**Figure 6:**
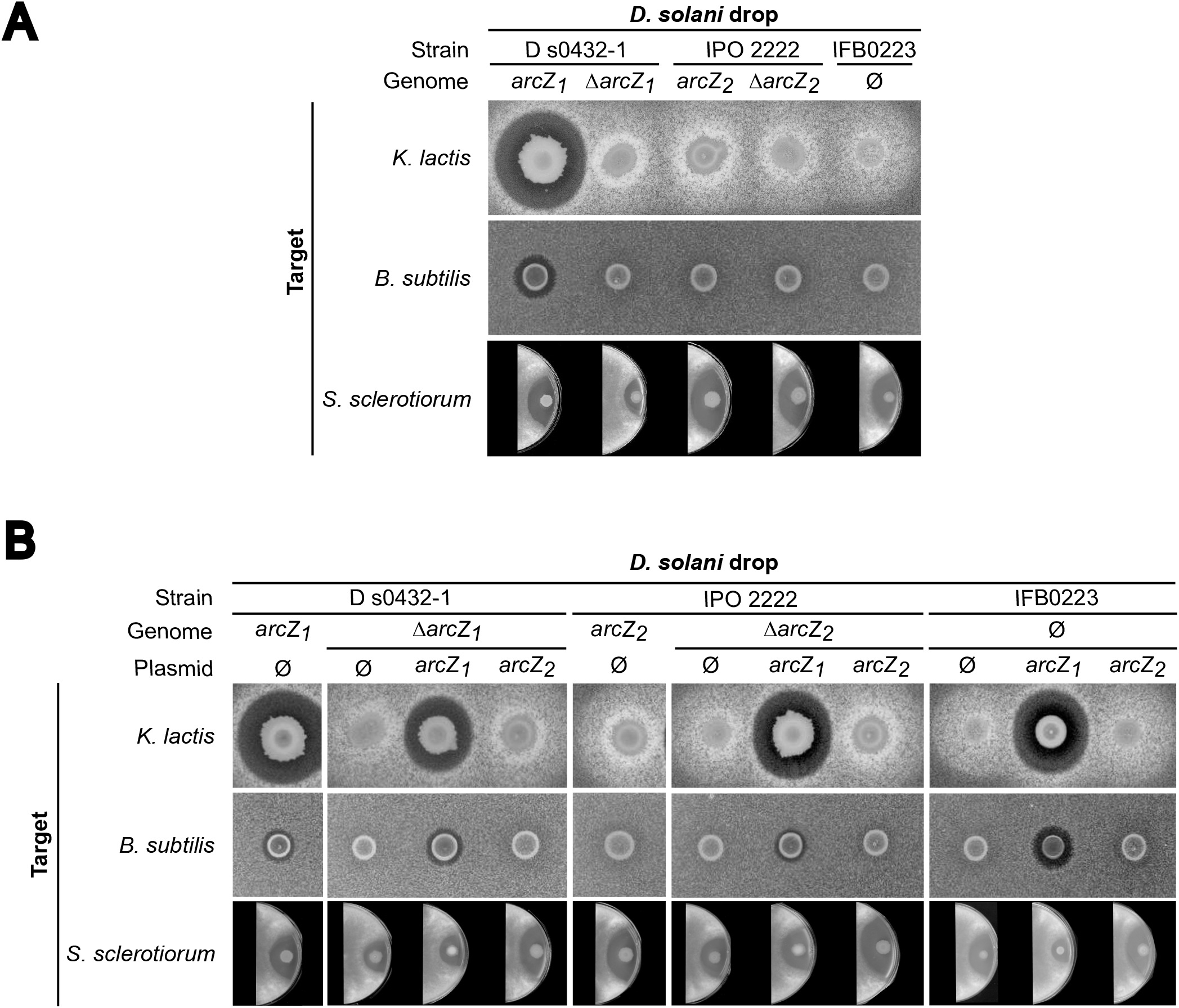
Inhibition of *K. lactis* and *B. subtilis* by *D. solani* is regulated by ArcZ. Yeast bioassay plates were prepared by mixing culture of *K. lactis* with warm YPD agar. Bacteria bioassay plates were prepared by mixing culture of *B. subtilis* with warm LB agar. PDA plates were inoculated with plugs of *S. sclerotiorum*. For each experiment, 5 µL of bacterial culture at OD_600nm_=2 from the different *D. solani* strains were spotted onto the plate. (A) Inhibition zones observed with Ds0432-1, IPO 2222, their respective *ΔarcZ* mutants and IFBO223. ArcZ_1_ is required for the synthesis of metabolites produced by the clusters *ssm* and *zms* (inhibition of *K. lactis* and *B. subtilis*, respectively) but not for the metabolite produced by the cluster *ooc* (inhibition of *S. sclerotiorum*). (B) Complementation tests and heterologous expression of the alleles *arcZ*1 or *arcZ*2 in the three *D. solani* strains. ArcZ_1_ activates the anti-microbial activities of *ssm* and *zms* clusters in IPO 2222 and IFBO223. All experiments were carried in 4 replicates.

To verify the effect of the G90A mutation in *arcZ*, we performed a heterologous complementation experiment by transferring the allele *arcZ*_*1*_ from D s0432-1 or *arcZ*_*2*_ from IPO 2222 in *D. solani* WT strains D s0432-1 or IPO 2222 or their Δ*arcZ* derivatives. The alleles *arcZ*_*1*_ and *arcZ*_*2*_ were also transferred into *D. solani* IFB0223 naturally deleted for *arcZ*.

### The G90A mutation in arcZ_*2*_ of *D. solani* IPO 2222 causes a loss of function of *zms* and *ssm* clusters

The two allelic forms *arcZ*_*1*_ and *arcZ*_*2*_ were cloned with their own promoter in the plasmid pWSK29-oriT, a mobilizable low copy plasmid with pSC101 origin and ampicillin resistance gene. DNA sequencing confirmed that the only difference between the two plasmids is the G90A mutation in pWSK29-oriT-*arcZ*_2_. The recombinant plasmids and the empty vector were transferred by conjugation into WT or Δ*arcZ D. solani* D s0432-1 or IPO 2222 strains, as well as WT *D. solani* IFB0223 (naturally deleted for *arcZ*). The phenotypes of the conjugants were determined by testing their anti-microbial activity against *B. subtilis, K. lactis* and *S. sclerotiorum* (Fig. 6B).

The transfer of *arcZ*_*1*_ in Δ*arcZ*_*1*_ Ds0432-1 restored the WT phenotypes, proving that the allele *arcZ*_*1*_ activates the anti-microbial activities of the *ssm* and *zms* clusters, while the anti-fungal activity was unaffected. In contrast, the transfer of *arcZ*_*2*_ in the Δ*arcZ*_*1*_ DS0432-1 strain did not restore the WT phenotypes. Thus, the difference in complementation between *arcZ*_*1*_ and *arcZ*_*2*_ is due to the presence of the G90A mutation that impairs ArcZ_2_ function.

An identical result was obtained with the Δ*arcZ*_*2*_ mutant of IPO 2222 or the IFB0223 strain carrying the pWSK29-oriT, pWSK29-oriT-*arcZ*_*1*_ and pWSK29-oriT-*arcZ*_*2*_. Indeed, the transfer of *arcZ*_*1*_ in strains WT IPO 2222, Δ*arcZ*_*2*_ IPO 2222 or IFB0223 activates the anti-microbial activities of the *ssm* and *zms* clusters. In contrast, the transfer of *arcZ*_*2*_ in these strains did not activate the *ssm* and *zms* anti-microbial activities. These data demonstrated that the presence of *arcZ* and the nature of the *arcZ* allele are the main factors governing the difference in phenotypes for antimicrobial activities between the *D. solani* strains D s0432-1 and IPO 2222 or IFB0223.

Finally, we questioned if an *arcZ* mutation could be found in additional *D. solani* natural isolates. We extended this genomic analysis to 57 *D. solani* genomes, including IPO 2222. It revealed that only four strains have an *arcZ* single-nucleotide mutation, all at different positions in the 3’ region of *arcZ*. It is not known whether these other mutations affect the antimicrobial activity of these strains. The *arcZ* DNA sequence of *D. solani* IPO 2222 was also used as a query in a BlastN search against the NCBI nucleotide database. Three *D. fangzhongdai strains* DSM 101947 (CP025003.1), QZH3 (CP031507.1), and LN1 (CP031505.1) had the G90A mutation in *arcZ*. A G90T mutation was also discovered in *D. parazeae* Ech586 (CP001836.1) (Fig. S5). Therefore, the G90 mutation to A or T of *arcZ* can be found in other *Dickeya* species.

### The G90A mutation in *arcZ*_2_ of *D. solani* IPO 2222 alters ArcZ_2_ processing

To gain further insight into the consequence of the G90A mutation in *arcZ*, Northern-blot experiments were performed to detect the sRNAs ArcZ_1_ in WT D s0432-1 and ArcZ_2_ in WT IPO 2222 (Fig. 7). A probe specific to the 3’ part of ArcZ_1_ and ArcZ_2_ was used (annealing from nucleotide 91 to 132). An abundant smaller transcript (~60nt) representing the processed 3′ fragment of ArcZ_1_ was detected in WT D s0432-1 but not in the mutant D s0432-1 Δ*arcZ*_1_ (Fig. 7, compared lanes 1 and 2). The full-length form corresponding to the ArcZ_1_ precursor was barely detected. This confirms that the processed form of ArcZ_1_ is predominant, like observed in *E. coli, Salmonella*, and *Photorhabdus* [38–40]. In IPO 2222, only the full-length ArcZ_2_ was detected in the WT strain whereas no ArcZ_2_ transcript was detected in the IPO 2222 Δ*arcZ*_*2*_ mutant (Fig. 7, compared lanes 5 and 6). This result suggests that the G90A mutation present in the allele *arcZ*_*2*_ of IPO 2222 prevents ArcZ_2_ processing in a shorter functional transcript. Next, we wondered if we could relate the importance of ArcZ processing to the activation of the *ssm* and *zms* clusters. To do that, Northern-blot analyses were realized with the *D. solani ΔarcZ* strains expressing either the *arcZ*_*1*_ or *arcZ*_*2*_ allele from the plasmid pWSK29-oriT-*arcZ*. On one hand, Northern blot showed that D s0432-1Δ*arcZ*_*1*_, IPO 2222 Δ*arcZ*_*2*_ and IFB0223 (natural *ΔarcZ*) expressing *arcZ*_*1*_ from D s0432-1 accumulate the processed short transcript of ArcZ_1_ (Fig. 7, lanes 3, 7 and 10) and have the capacity to inhibit yeast and bacteria (Fig. 6). On the other hand, D s0432-1Δ*arcZ*_*1*_, IPO 2222 Δ*arcZ*_*2*_ and IFB0223 expressing *arcZ*_*2*_ from IPO 2222 showed the accumulation of the non-processed transcript of ArcZ_2_ (Fig. 7, lanes 3, 7 and 10) and an inability to inhibit yeast and bacteria (Fig. 6). These results supported the hypothesis that the G90A mutation in ArcZ alters the processing of the ArcZ precursor in a shorter form, leading to its inability to play a correct regulatory function on the *ssm* and *zms* clusters.

**Figure 7.**
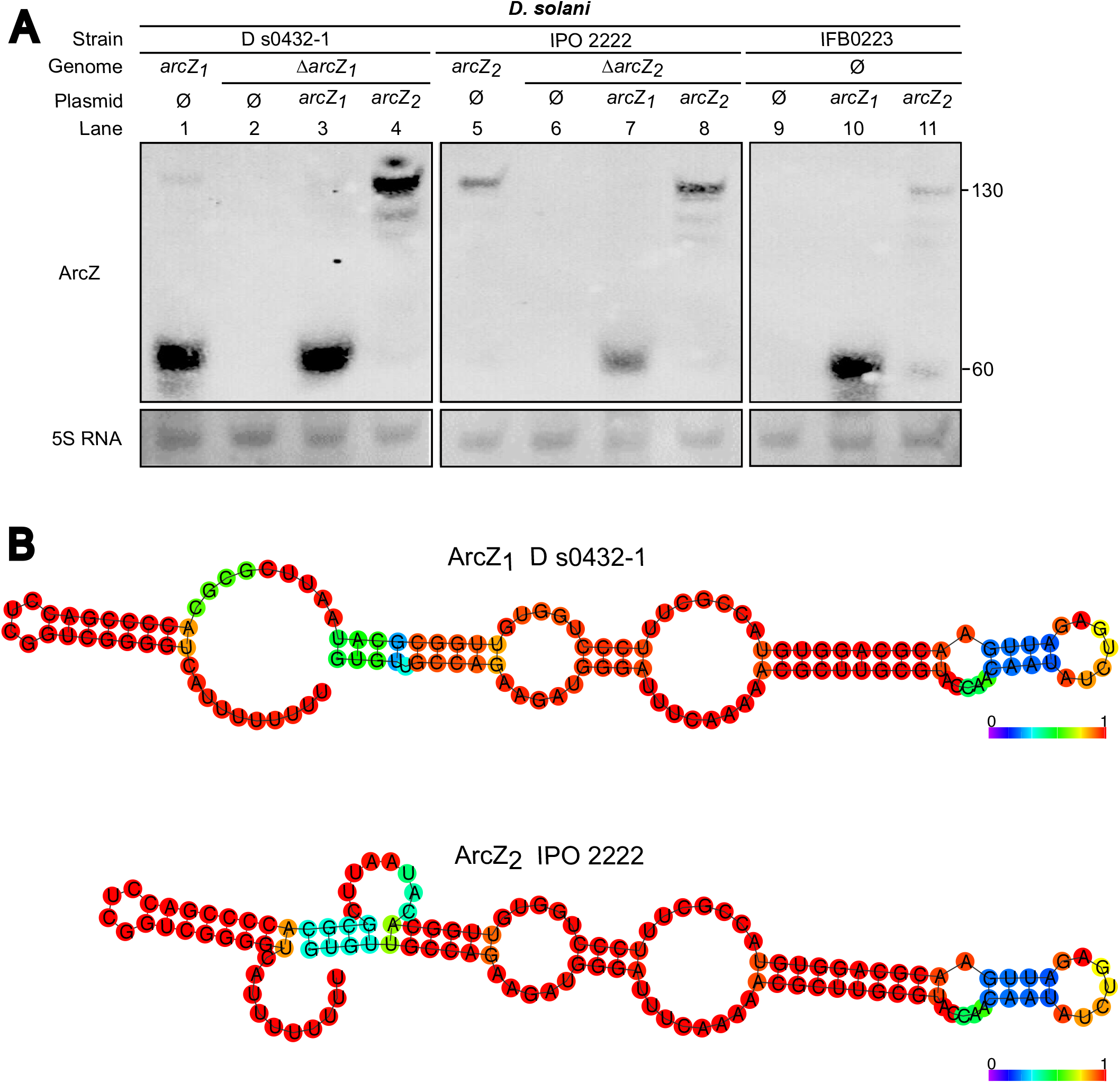
Expression of different ArcZ alleles: the G90A mutation in ArcZ_2_ does not prevent its production but its processing. (A) Detection of the sRNA ArcZ_1_ and ArcZ_2_ by Northern Blot in *D. solani* strains. A small transcript (~60 nt) corresponding to the processed 3′ fragment of ArcZ_1_ was detected in D s0432-1 and in all strains expressiong *arcZ*_*1*_ on plasmid. In IPO 2222, only the full-length form (~132nt) ArcZ_2_ was detected, as well as in strains expressing the allele *arcZ*_*2*_ on plasmid. The 5S RNA is used as a loading control. This experiment was performed twice with biological replicates and gave identical results. (B) Prediction of secondary structure of the precursor forms of ArcZ_1_ and ArcZ_2_ by RNA fold. Bases are colored according to base-pairing probabilities (see color scale). The G90A mutation present in ArcZ_2_ seems to modify the structure of the ArcZ sRNA precursor.

## DISCUSSION

PKS/NRPS secondary metabolite pathways are a great source of molecules with anti-eukaryotic or anti-prokaryotic activity, giving the bacteria that synthesize them a competitive advantage over other organisms. Targeted mutagenesis of the *D. solani* D s0432-1 chromosome has allowed us to specifically study the involvement of three secondary metabolite biosynthesis clusters encoded by all the *D. solani* strains for which genome sequence is available. We focused our work on strain D s0432-1, one of the most virulent *D. solani* strains [18]. We have demonstrated that *D. solani* D s0432-1 is able to inhibit the growth of a variety of living microorganisms. We tested bio-activities against Gram-negative bacteria (*E. coli, P. atrosepticum, D. dadantii*), Gram-positive bacteria (*B. subtilis, S. scabiei*), yeasts (*S. cerevisiae, K. lactis, C. albicans*), and fungi (*B. cinerea, M. oryzae* and *S. sclerotiorum*). The cluster *ooc* encodes a biosynthesis pathway that produces an oocydin-like molecule, a chlorinated macrocyclic lactone molecule having antifungal, antioomycete and antitumor activities in *S. plymuthica* [23,29]. It was previously shown that the *D. solani* strains MK10, MK16 and IPO 2222, all encoding the oocydin cluster, inhibit the growth of the fungus *V. dahliae* and the oomycete *P. ultimum* [29]. Our study showed that *D. solani* D s0432-1 prevents the growth of the fungi *B. cinerea, M. oryzae* and *S. sclerotiorum*, and that inactivation of the *oocL* gene suppressed the inhibition of the three fungi without any visible effect on the bacteria or yeasts tested.

By testing other mutants of the other secondary metabolite clusters, we showed that the mutant Δ*ssmG* was nearly completely unable of preventing the growth of *S. cerevisiae* and *K. lactis*. Therefore, the *ssm* cluster appears to be responsible for yeast inhibition. *D. solani* D s0432-1 also inhibits the human pathogen yeast *C. albicans*, but its inhibitory activity appears to be weaker than that observed with *S. cerevisiae* and *K. lactis* in our inhibition assay. This observation on the *ssm* cluster paves the way for the identification of a new molecule with highly specific inhibitory activity against yeasts. The *ssm* cluster is conserved in several *Dickeya* genomes but rarely in other bacterial genera, except in some *Rouxiella* species. The structure of the molecule and its target remains to be elucidated.

Our study also showed that *D. solani* D s0432-1 has an anti-bacterial activity linked to the cluster *zms*, which encodes a zeamine biosynthetic pathway. Zeamine produced by *S. plymuthica* kills *B. subtilis*, the yeasts *S. cerevisiae* and *Schizosaccharomyces pombe*, and the nematode *Caenorhabditis elegans* [32]. We showed that *D. solani* D s0432-1, but not the Δ*zmsA* mutant, can clearly inhibit the growth of two Gram-positive bacteria, *B. subtilis* and *S. scabies*. Since *D. solani* and *S. scabiei* are two potato pathogens, they might be in competition for the same ecological niche. It is thus interesting to note that *D. solani* can inhibit the growth of another potato pathogen. We also observed a slight inhibition exerted by *D. solani* D s0432-1 against *E. coli*, but no inhibition towards *D. dadantii* and *P. atrosepticum*. Zeamine resistance of *D. dadantii* could be explained by the presence in its genome of the genes *desAB* encoding an RND pump identified in *D. oryzae* EC1 and involved in zeamine efflux [41]. In-frame deletion of *desA* or *desB* in *D. oryzae* EC1 leads to a zeamine sensitive phenotype [41]. In *in planta* Tn-seq experiments with *D. dadantii* 3937 [42], mutants of the genes *desA* and *desB* (Dda3937_00787 and Dda3937_00786 respectively) did not display a significant negative or positive variation (Log2 fold-changes −0.09 and +0.44, respectively), suggesting that this RND efflux pump does not play a significant role during *D. dadantii* chicory infection, contrary to the *D. dadantii* RND efflux pump AcrAB that appeared to be essential for virulence [42]. While *P. atrosepticum* is not inhibited by *D. solani* D s0432-1, it does not have the genes *desAB*. Zeamine resistance could be provided by another efflux pump or by a different mechanism in *P. atrosepticum*. It’s worth remembering that the *D. solani* D s0432-1 Δ*zmsA* mutant grows slightly faster than the WT strain after reaching OD_600nm_ 0.4 (Fig. S1). This small difference in growth might be explained by the lack of need of the mutant to adapt to the presence of zeamine in the medium. We also observed that *D. solani* D s0432-1 Δ*ssmG* produced a very slight halo of inhibition against *S. cerevisiae* and *K. lactis*, barely visible in our tests. This weak halo disappears in the double mutant Δ*ssmG*Δ*zmsA* and the triple mutant *ΔssmGΔoocLΔzmsA*, showing that the zeamine produced by the *D. solani zms* cluster has also an anti-yeast action. It is worthwhile to remind that zeamine produced by *S. plymuthica* A153 is bioactive against *S. cerevisiae* and *S. pombe* [32].

Thus, the presence of three clusters encoding secondary metabolites allows *D. solani* D s0432-1 to produce an arsenal of bioactive secondary metabolites against a variety of living microorganisms. The cluster *ssm* produces an unkown molecule active against yeasts; the cluster *ooc* produces an oocydin-like molecule active against fungi and the cluster *zms* produces a zeamine-like molecule active against bacteria and yeast. In future studies, It would be instructive to investigate *D. solani* behavior toward other organisms found in the environment, such as amoebas, nematodes, or even insects.

We then expanded our studies to a larger set of *D. solani* strains isolated in different European countries and years. Through the combination of phenotypic analysis and comparative genomics, confirmed by mutant construction, we demonstrated that the ArcZ sRNA plays a major role in activating the antimicrobial activities of the clusters *ssm* and *zms*. We first observed that some *D. solani* WT strains are defective for yeast inhibition and possess different *arcZ* alleles, a deletion in the case of strain IFB0223, or the presence of a SNP G90A in the 3’ part of *arcZ* in the type strain IPO 2222. We demonstrated that these arcZ alterations significantly impair the *D. solani* ability to inhibit other bacteria and yeasts. These *arcZ*-related genomic changes represent a minority of *D. solani* isolates, only two of the nine strains phenotypically tested. Hence, our results show that despite a probable clonal origin, environmental strains of *D. solani* can present very different characteristics due to a single mutation into a regulatory sRNA. The *arcZ* polymorphism may be one of the explanations for the variability of phenotypes and virulence observed between genetically close *D. solani* strains. A global investigation of the polymorphism of all the short RNAs within the genomes of the same species could give surprising results. This study shows that it is critical to pay attention to the allelic diversity due to SNPs present not only in protein coding genes but also in sRNA genes.

ArcZ is a well-known regulatory sRNA that is associated with the chaperone Hfq. The ArcZ primary transcript (121 to 134-nt) is converted by the essential endoribonuclease RNAse E [43] to a shorter stable approximative 60-nt RNA retaining the seed region. The precise length of this short processed RNAs depends on the bacterial species [38,40,44,45]. In *E. coli*, the processed short form of ArcZ promotes *rpoS* translation while suppressing the expression of several other genes. In anaerobic conditions, *arcZ* is repressed and translation of RpoS is low [44]. RpoS is the RNA polymerase alternative sigma factor that governs the general stress response, which is activated by a wide range of conditions. It controls 10% of the *E. coli* genome [46]. RpoS can control positively or negatively the production of metabolites such as antibiotics [29,47]. *rpoS* mRNA is not the only ArcZ direct target. In *Photorhabdus* and *Xenorhabdus*, ArcZ pairs with the mRNA encoding HexA, a transcriptional repressor of the expression of specific metabolite gene clusters [40]. In *Dickeya* species, the regulator gene *pecT* is the orthologous gene to *hexA*. In *D. dadantii* 3937, the *pecT* mRNA is also a direct target of ArcZ. Short processed ArcZ binds to Hfq and directly to the 5’UTR region of *pecT* mRNA, repressing its translation [45]. PecT is directly and indirectly involved in the expression of genes encoding plant cell wall degrading enzymes and virulence factors [45]. In the phytopathogen *Erwinia amylovora*, ArcZ participates also in the positive control of T3SS, exopolysaccharide production, biofilm formation, and motility [48]. Finally, another known direct target of ArcZ is the *flhD* mRNA [49,50]. FlhD forms with FlhC the master regulator FlhDC that controls motility in bacteria [51]. Although we demonstrated the critical role of ArcZ in the regulation of secondary metabolite production in *D. solani*, it remains to understand the precise mechanism of ArcZ action on the ssm or zms clusters. We can only speculate that this could occur through the control of *rpoS, flhD*, or *pecT*, three regulators known to be ArcZ targets. However, other actors could be involved. Another point to consider is a more global role of *D. solani* ArcZ exerted on other cellular functions such as motility and virulence. ArcZ targets in *D. solani* need to be investigated in the future. It is worth pointing out that the G90 nucleotide is in the 3’ region of ArcZ that interacts with the target mRNAs *rpoS, flhD*, or *pecT* in other bacteria (Fig. S6). Northern-blot analysis revealed that ArcZ_2_ of *D. solani* IPO 2222 is not processed in the stable shorter sRNA as ArcZ_1_ of *D. solani* D s0432-1. The G90A mutation in IPO 2222 ArcZ_2_ is predicted to modify the secondary structure of the ArcZ precursor form (Fig. 7B). Since maturation by RNase E is essential for target regulation by the ArcZ sRNA in *Salmonella*, we conclude that the absence of a short form of ArcZ in IPO 2222 is probably the main cause of the loss of regulatory function of the IPO 2222 ArcZ_2_.

The type strain *D. solani* IPO 2222 is largely used in academic laboratories as a model to study the *D. solani* species. In light of our findings, it could be important to compare data obtained with the strain IPO 2222 with those obtained with other *D. solani* strains. The *arcZ* mutations may be responsible, at least partially, for the lower virulence of IPO 2222 and IFB0223 observed previously in comparison to D s0432-1 [18]. Even strains with the same *arcZ, ssm, ooc*, and *zms* genomic sequences as D s0432-1, showed heterogeneity in their ability to inhibit other micro-organisms. For example, IPO 3494 and IFB0484 inhibited *B. subtilis* better than other *D. solani* strains (Fig. 5A). Other specific mutations in the genomes of these strains, which could also modify the activity of secondary metabolite clusters, could explain these phenotypic variations.

We first described the inhibitory function of the *ssm* cluster of the *D. solani* strain D s0432-1 against the yeasts *S. cerevisiae* and *K. lactis* in a 2021 bioRxiv preprint [27]. Very recently, in November 2022, Matilla *et al*. published results on the *ssm* cluster of the *D. solani* strain MK10 [26]. The *ssm* cluster was named *sol* in this study and the authors described the antifungal activity of the uncharacterized molecule that they called solanimycin [26]. Matilla *et al*. also report the inhibition observed with the *D. solani* strain IPO 2222 against *Verticillium dahliae* and *Schizosaccharomyces pombe* ([26], Fig S1). In our conditions, strain IPO 2222 was able to inhibit the fungus *Sclerotinia sclerotiorum* but not the yeast *Kluyveromyces lactis* (Fig. 5). In comparison to our study, Matilla *et al* used different fungal species but also different growth conditions (medium, temperature, exposure time, etc). By random mutagenesis of strain MK10, they identified RsmA and the two *Dickeya* quorum-sensing systems ExpI/R and Vfm as regulators of the *ssm/sol* cluster such [26]. Interestingly, the Rsm regulatory pathway of *D. dadantii* 3937 is regulated by PecT, which is under the ArcZ control [45]. Further studies are now necessary to specify the biological targets of solanimycin and to clarify the role in *ssm/sol* expression of the sRNA ArcZ, quorum-sensing, and transcriptional regulators PecT or RsmA, in order to better understand the regulatory network controlling the production of secondary metabolites in *D. solani*. Our lab is currently investigating the regulatory pathway of secondary metabolite clusters from ArcZ.

In conclusion, our results provide the basis for further investigation of a novel metabolite capable of limiting yeast growth and, at the same time, open doors to understand how the ArcZ sRNA acts as a global regulator to regulate two different clusters of secondary metabolites in *D. solani*. It also shows how a single mutation in a small RNA can greatly modify the phenotype of bacterial strains that are very close phylogenetically.

## MATERIAL AND METHODS

### Bacterial and fungal strains, plasmids and growth conditions

The *E. coli* and *Dickeya* bacterial strains, plasmids and oligonucleotides used in this study are described in Table S3 and S4. The genome accession number of *D. solani* D s0432-1 is NZ_CP017453. The following strains have been used in the study: *Sclerotinia sclerotiorum* S5, *Botrytis cinerea* B05.10, *Magnaporthe oryzae* Guy11, *Saccharomyces cerevisiae* BY4743 (*MATa/α his3Δ1/his3Δ1 leu2Δ0/leu2Δ0 LYS2/lys2Δ0 met15Δ0/MET15 ura3Δ0/ura3Δ0), Kluyveromyces lactis* MWL9S1 [52], *Candida albicans* SC5314, *Dickeya dadantii* 3937, *Pectobacterium atrosepticum* SCRI1043, *Streptomyces scabiei* CFBP4517, *Bacillus subtilis* PY79. *E. coli* was grown routinely at 37°C in LB. Fungus strains were grown at 25°C onto Potato Dextrose Agar (PDA). *S. scabiei* was grown in tryptic soy broth (TSB) medium at 28°C. *B. subtilis, P. atrosepticum* and the *Dickeya* strains were cultivated in LB unless specified. Yeast cells were grown at 30°C in rich medium consisting of complete yeast extract-peptone (YP) medium containing 1% Bacto yeast extract, 1% Bacto peptone (Difco) supplemented with 2% glucose (yeast extract-peptone-dextrose [YPD] medium). For the bacteria, yeast and fungi inhibition assay, M63 medium supplemented with sucrose (2 g (NH_4_)_2_SO_4_, 13.6 g KH_2_PO_4_, 2.5 mg FeSO_4_7H_2_O, 0.2 g MgSO_4_7H_2_O, 10 g sucrose, per liter) were employed to grow overnight the *D. solani* strains before performing the assay.

When required, antibiotics were added at the following concentrations: ampicillin (Amp), 100 µg/L; nalidixic acid (Nal), 10 µg/mL. Diaminopimelic acid (DAP) (57 µg/mL) was added for the growth of the *E. coli* MFDpir strain. Media were solidified with 12 g/L agar.

### Growth inhibition assay of bacteria and yeast

*D. solani* strains were grown for 24 h at 30°C with shaking in M63 medium supplemented with 1% sucrose. *B. subtilis, E. coli, D. dadantii* and *P. atrosepticum* were grown in LB at 30°C with shaking overnight. The next day, the OD_600_ of the cultures of *D. solani* were adjusted to 2. The temperature of melted LB agar was lowered to around 40°C (just before the agar re-solidified). 100 mL of the LB agar in surfusion were mixed with 100 µl of the OD_600_ 1 culture of *B. subtilis, E. coli, D. dadantii* or *P. atrosepticum*. 30 mL of inoculated LB agar were poured in 12-by 12-cm square plates. Then 5 µl of the OD_600_ 2 cultures of *D. solani* were spotted onto the inoculated square plates which were incubated at 30°C for 24-48 h before visualization of the inhibition zone.

The same experiment was conducted with *S. scabiei*, except that it was grown for 3 days in TSB at 28°C and TSB agar was poured in the square plates. With the yeasts *S. cerevisiae, K. lactis*, and *C. albicans*, a similar protocol was used with YPD medium and with OD_600_ 2 culture of yeast per 30mL of medium.

### Growth inhibition of fungal strains

*S. sclerotiorum, B. cinerea* and *M. oryzae* were grown onto PDA plates at 25°C for 5, 7 and 10 days respectively. *D. solani* strains were grown for 24 h at 30°C with shaking in M63 medium supplemented with 1% sucrose. The OD_600_ of these overnight cultures were adjusted to 2. Then, 5 µl of the bacterial suspensions were spotted onto PDA plates with 5 mm agar plugs of fungus at the center of the Petri dish. The radius of fungus inhibition zone was measured.

### Construction of the MLSA tree positioning strains within the *Dickeya* genus

Details on the methods used for the construction of MLSA tree positioning strains are given in Supplementary Material and Methods.

### Constructions of the mutant strains and plasmids

All the material and methods for the construction of the strains and plasmids used in this study are given in Supplementary Material and Methods.

### RNA isolation and Northern detection

All the material and methods for RNA isolation and Northern detection performed in this study are given in Supplementary Material and Methods.

## ACKNOWLEDGMENTS

B.T. was supported by a PhD grant from the Ministère de l’Enseignement Supérieur, de la Recherche et de l’Innovation. We thank Veronique Utzinger for technical assistance, the members of the Membrane Trafficking and Signaling in Bacteria (MTSB) team for discussions, Amélie De Vallée, Nathalie Poussereau and Christophe Bruel for advices on fungi growth and providing the strains, Alexandre Soulard, Marc Lemaire and Jade Ravent for advices on yeast growth and providing the strains and medium. We would also like to thank the past reviewers whose comments allowed us to greatly improve this study.

## AUTHOR CONTRIBUTION

B.T., E.G. B.J. A.L., G.C., C.J., G.E carried out the experiments. G.E wrote the manuscript with support from B.T, E.G., A.L., R.M., F.D. and H-C-P.N.

F.D., C.J. H-C-P.N. and R.M. performed the phylogenetic and bioinformatic analyses. G.E conceived the original idea with advises from H-C-P.N. G.E supervised the project. All authors provided critical feedback and helped shape the research, analysis and manuscript.

## CONFLICT OF INTEREST

The authors state that the research was conducted in the absence of any commercial or financial relationship that could be interpreted as a potential conflict of interest.

## FUNDING

This work was supported by a grant from Agence Nationale de la Recherche to L.A. (Project RNAchap, ANR-17-CE11-0009-01) and to E.G. (Project Tn-Phyto, ANR-19-CE35-0016). E.G was also supported by the FR BioEEnVis, annual credits from the University Lyon I and the CNRS at regular basis.

## FIGURE LEGENDS

**Figure S1.**
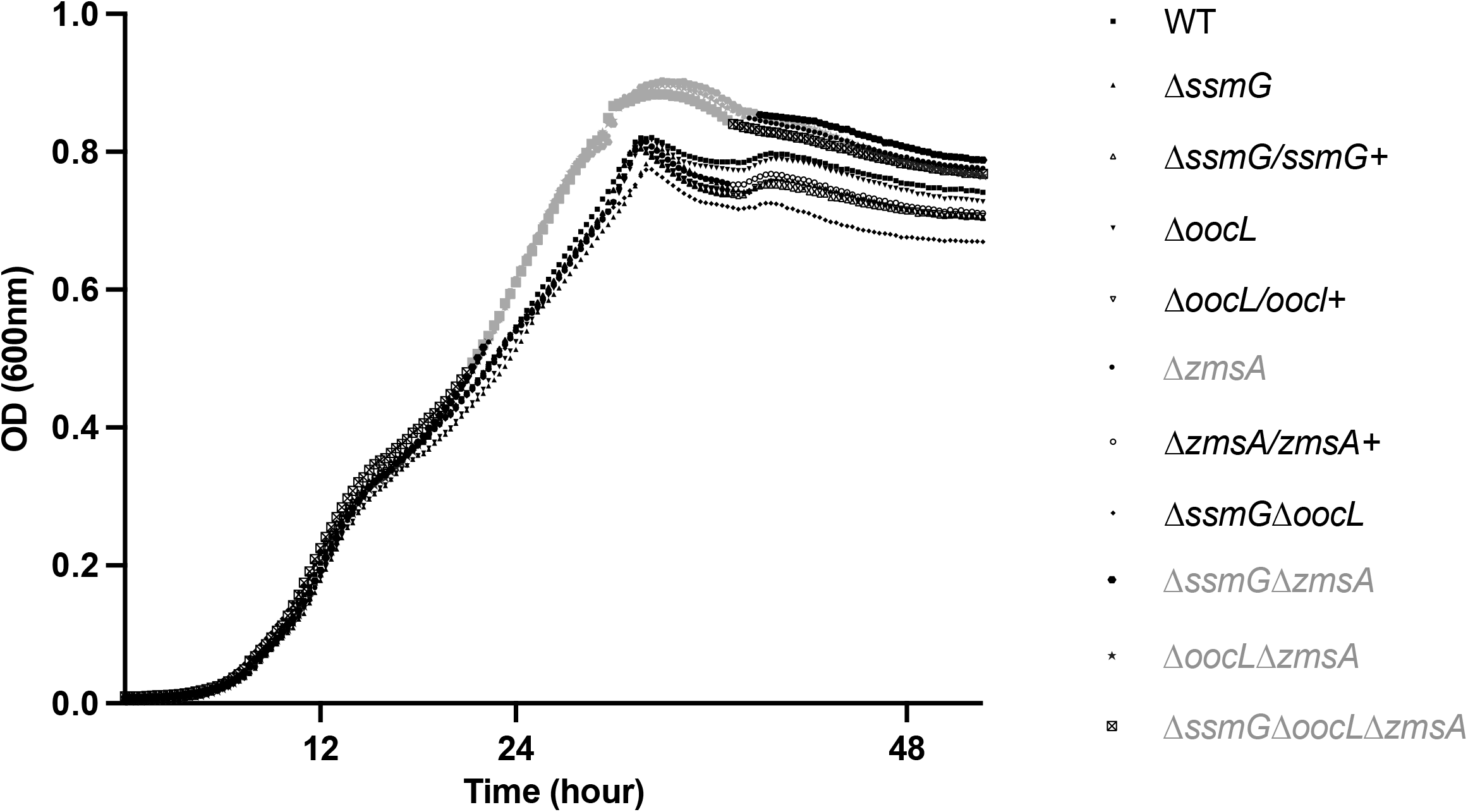
Growth curves of *D. solani* strains and mutant derivatives. A 96-well plate containing M63 medium with 1% sucrose was inoculated with the strains used in this study at an OD of 0.06. The growth of each strain was determined by measuring OD_600_ every 20 minutes during 2 days in TECAN. The different strains showed similar overall growth, except for the light grey points where a low but significant fitness gain was observed for the mutants *Δzms, Δssm Δzms, Δooc Δzms* and Δ*ssm Δooc Δzms* (Mann-Whitney test, p-value<0.05).

**Figure S2.**
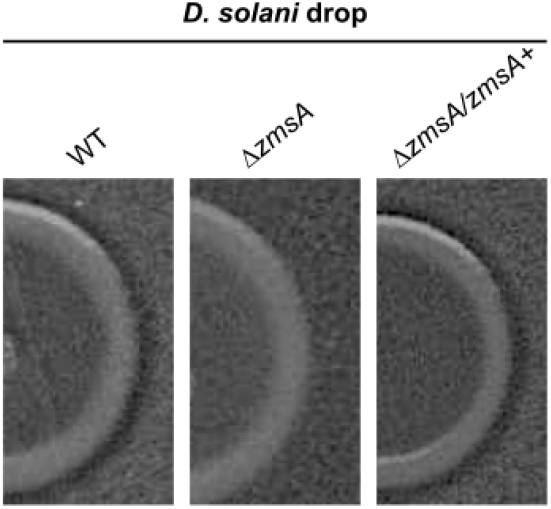
Inhibition of *E. coli* growth by *D. solani* D s0432-1 and mutant derivatives (zoom on drop borders). Bioassay plates were prepared by mixing *E. coli* culture with melted LB agar as described in experimental procedures. 5 µL of bacterial culture at OD_600nm_ = 2 of *D. solani* D s0432-1 (WT) or derivatives were spotted onto the plate and incubated at 30°C during 48 h. A slight inhibition zone was observed except with the *Δzms* mutant. All experiments were carried in 4 replicates.

**Figure S3.**
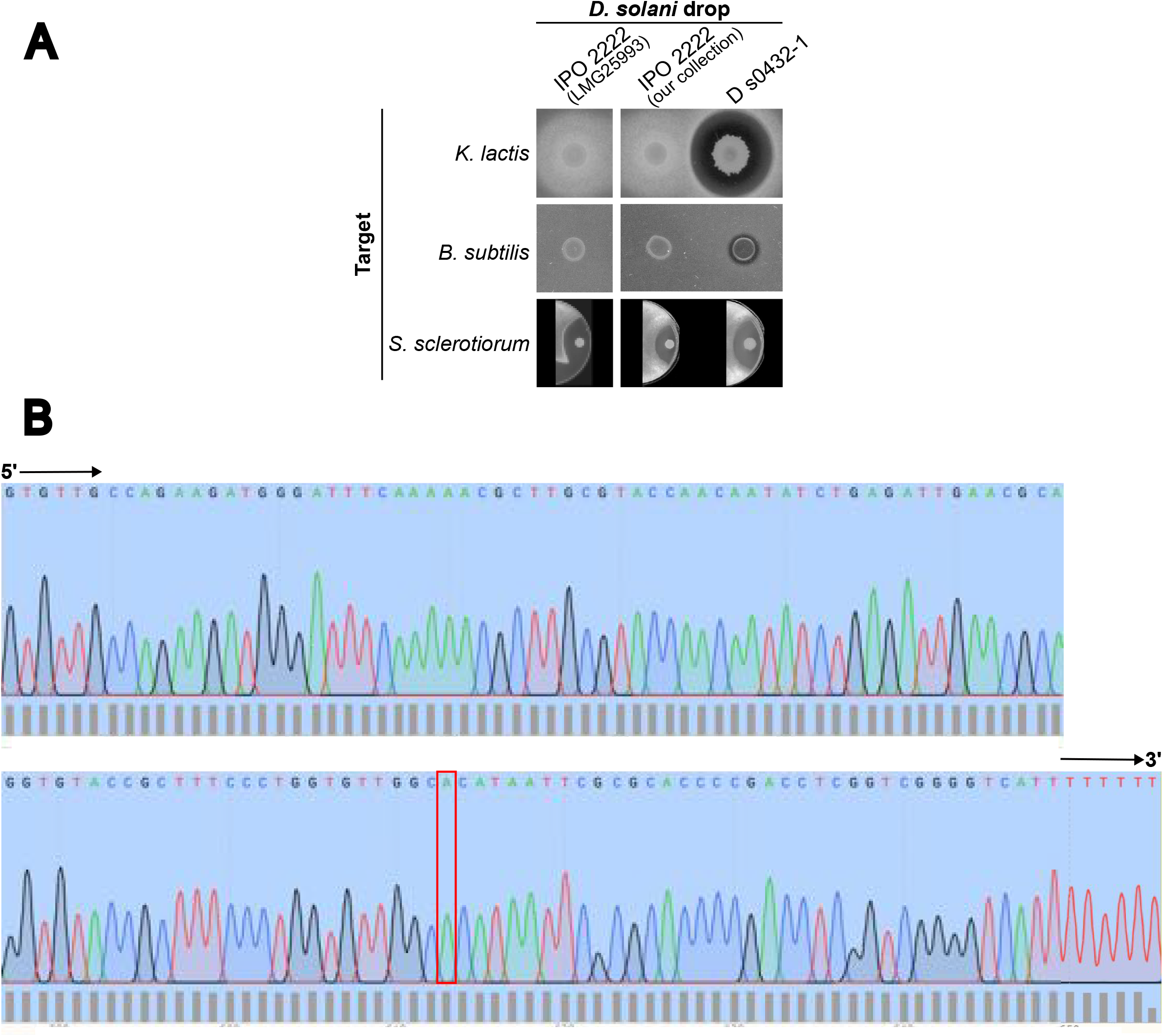
Phenotypes and *arcZ* sequence of WT *D. solani* IPO 2222 from the BCCM collection (LMG 25993). (A) Inhibition assay of *K. lactis, B. subtilis* and *S. sclerotiorum* by WT *D. solani* IPO 2222 LMG 25993 from BCCM collection, compared to the strains used in this study. (B) Sanger sequencing results of *D. solani* IPO 2222 *arcZ* (LMG 25993). The mutation G90A is highlighted by a red frame.

**Figure S4.**
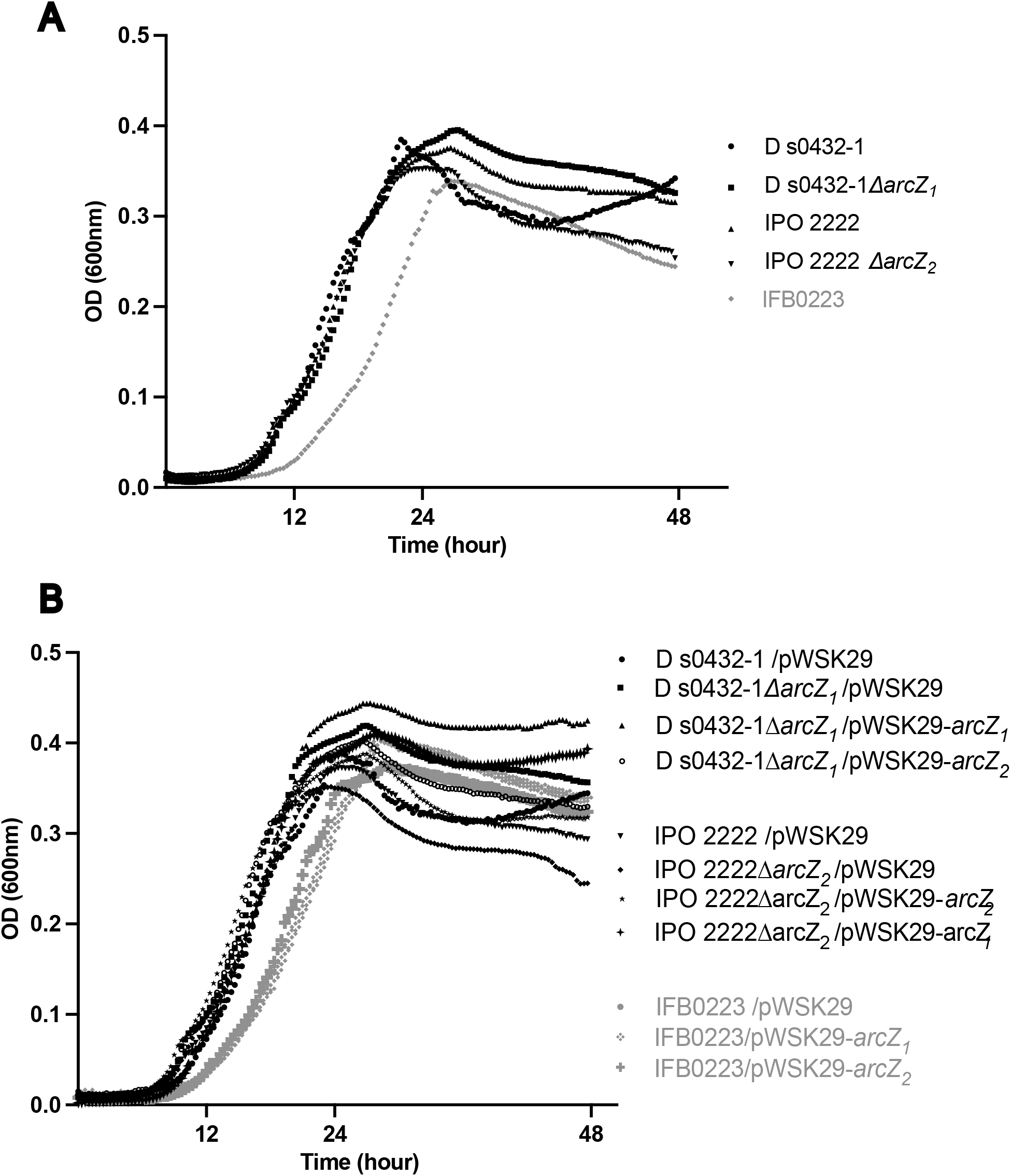
Growth curves of WT and Δ*arcZ D. solani* strains used in this study. (A) Growth curves of D s0432-1, IPO 2222, their respective *ΔarcZ* mutants and IFB0223. The latter (in grey) has a slight growth delay (Mann-Whitney test, p-value<0.05). (B) Growth curves of strains used in complementation and heterologous expression tests. The plasmids do not cause any growth defect in D s0432-1, IPO 2222 and *ΔarcZ* mutants. IFB0223 strains (in grey) have growth delay already observed in the experiment without the plasmid (Mann-Whitney test, p-value<0.05). All experiments were carried in 4 biological replicates.

**Figure S5.**
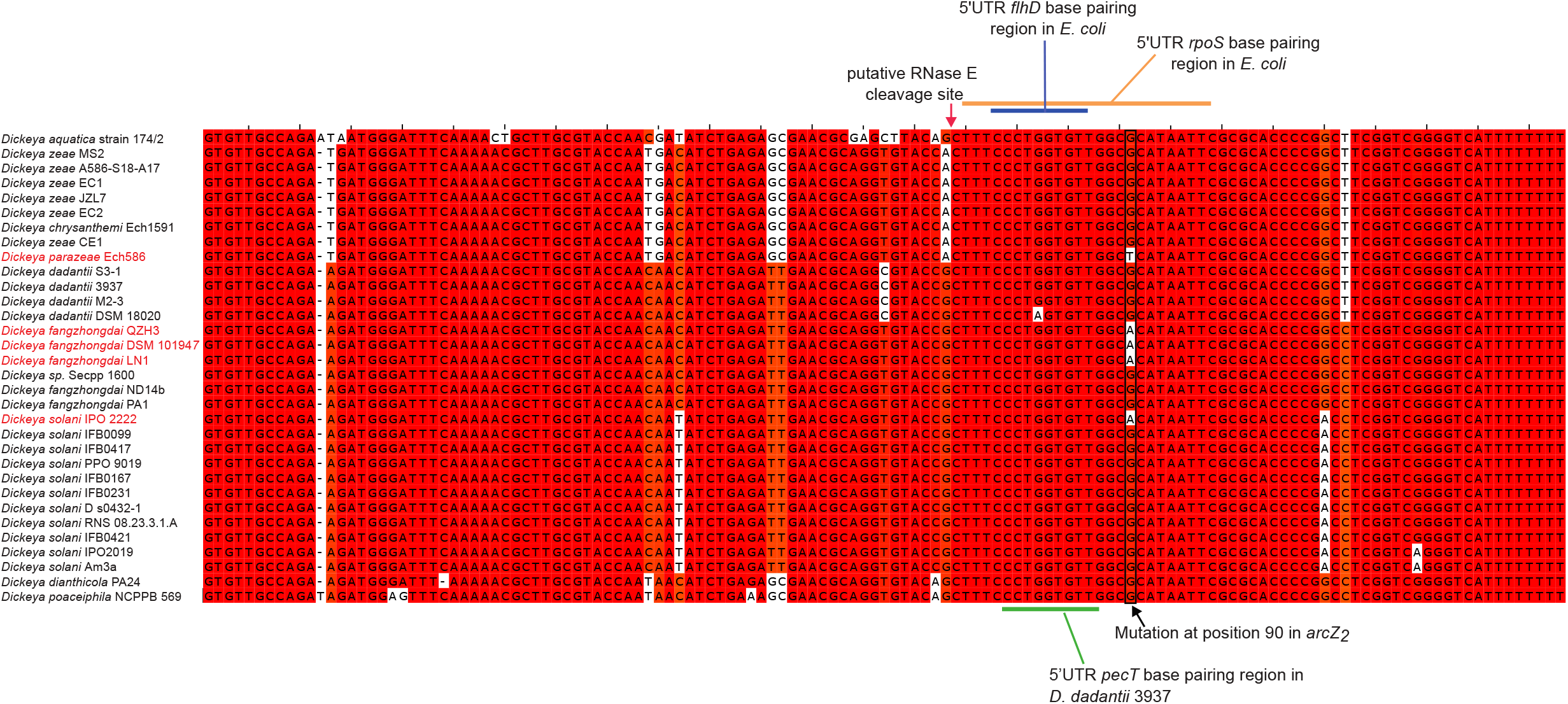
Alignment of *arcZ* sequences. The *arcZ* DNA sequences were retrieved by running a BlastN on the NCBI database using the *arcZ* sequence of *D. solani* D s0432-1 as query. Search was limited to the *Dickeya* genomes. Then, the *Dickeya arcZ* sequences were aligned with *E. coli* MG1655 *arcZ* by using Jalview [53] and Muscle [54]. Known regions of interactions with the *pecT, flhD* and *rpoS* 5’UTR mRNA in *D. dadantii* and *E. coli* are indicated

## Supplementary Material and Methods

DOCX file with supplementary material and methods.

**<u>Table S1</u>. Distribution of the clusters *ssm, ooc*, and *zms* in 155 *Dickeya* strains whose genome is sequenced**. For each *Dickeya* species, the strains were classified on the basis of the presence or absence of the cluster *ssm, ooc*, and *zms*.

**<u>Table S2</u>. Lists of SNPs in *D. solani* IFB0099, D S0432.1, IFB0223, IPO3337 and IPO 2222**.

**<u>Table S3</u>. Strains and plasmids used in the study**.

**<u>Table S4</u>. Oligonucleotides used in the study**.

## Notes

### Competing Interest Statement

The authors have declared no competing interest.

### Summary of Updates

Since the first version of the manuscript posted on BioRxiv where we revealed the discovery of a new secondary metabolite cluster in Dickeya solani that inhibits yeast growth, we made a discovery that forced us to postpone the submission of the manuscript to a journal: some wild type strains of D. solani are impaired in their ability to kill yeast and bacteria. Further investigations led us to the discovery that this loss-of-function phenotype is due to a single mutation in the small regulatory RNA ArcZ. The new version of this manuscript describes this finding.

