## Supplementary material for "A natural single nucleotide mutation in the small regulatory RNA ArcZ of *Dickeya solani* switches off the antimicrobial activities against yeast and bacteria": Supplementary Material and Methods.docx

**Supplementary Figure S6: supplementary material and methods**

**PCR conditions**

To amplify DNA for cloning, a bacterial suspension was prepared by boiling 10 minutes at 95°C an isolated colony of *D. solani* suspended into 40 µl of sterilized water. The PCR was carried out in a 50 µl reaction mix containing 1 µl of the bacterial suspension, 1.5 µl of 10 µM of each primer (Eurofins), 25 µL of Primestar master mix 2x (Takara) and demineralized water. Thermocycling program followed the manufacturer recommendation.

To check the correct deletion of a gene of interest in *D. solani*, a bacterial suspension was made as described before. The suspension was centrifuged 1 min at 12000 g to remove cellular debris. The supernatant was used as a template for PCR.

To amplify up to 3-kb DNA fragments, colony PCR was performed with DreamTaq Green PCR Master mix (2x) (Thermofisher, Waltham, MA, USA). For fragments larger than 3-kb, the colony PCR was carried out with the Q5 High-Fidelity DNA Polymerase (Biolabs) by following manufacturer recommendations.

**Construction of the Nal^R^ Gm^R^ derivative of *D. solani* D s0432-1**

A spontaneous mutant of *D. solani* D s0432-1 resistant to nalidixic acid (Nal^R^) was obtained by growing the WT strain at 30°C in LB medium supplemented with 5 µg/mL Nal for 18 h before spreading the liquid culture onto LB agar plates with Nal at 10 µg/mL. The Nal^R^ strain was named DS50 (Table S1). Then a Gm^R^ derivative of strain DS50 was constructed by insertion of a mini-Tn7-Gm cassette into the attTn7 site of *D. solani,* as previously performed with *D. dadantii* 3937 [1] by using the plasmids pTn7-M [2] and pTnS3 [3]. The correct integration of the Gm^R^ cassette was checked by colony PCR using oligo pairs L365/L848 (amplification of 350 bp for the correct integration) (Table S2). This gave the parental Nal^R^ Gm^R^ strain DS58. This modified strain of *D. solani* D s0432-1 has the same phenotype for anti-microbial activities as the genetically non-modified WT strain (data not shown). To simplify the understanding of the experiments, this strain is designated WT in figures 2, 3, 4, S1 to S4.

**Construction of single, double and triple in-frame deletion mutants of the *ssm*, *ooc* and *zms* clusters.**

To construct in-frame deletion mutants, the *sacB* counter-selection method was used [4]. pRE112 is an R6K-based suicide plasmid carrying the *sacB* gene and the *cat* gene (Cm^R^). Two PCR fragments corresponding to the upstream and downstream 0.5-kbp DNA of the gene to be deleted were cloned into SacI/KpnI digested pRE112 using the Gibson's assembly method. Chemical ultracompetent DH5α λpir cells were prepared with the Mix & Go! *E. coli* Transformation Kit using standard procedures (Zymo Research). Transformants were selected onto LB plate supplemented with chloramphenicol. Colonies with the correct plasmid were selected by colony PCR with oligo pairs L762/L763 and DreamTaq DNA polymerase (Thermofisher, Waltham, MA, USA). Plasmids were extracted with the NucleoSpin Plasmid kit (Macherey-Nagel) and checked by restriction digestion (NEB) and sequencing (Eurofins). Then, plasmids were transferred into competent *E. coli* strain MFDpir [5] prepared with the TSS method [6]. *E. coli* MFDpir produces the RP4 conjugation machinery, which allows the transfer of the suicide plasmid into *D. solani* by conjugation. For conjugation, a few colonies of *D. solani* and MFDpir were mixed in the same proportion in 500 µl LB and centrifuged for 2 min at 8000 rpm. The pellet was resuspended in 90 µl LB with 5 µl DAP at 57 mg/mL, and deposited onto a LB agar plate incubated at 30°C. After 18 h, the bacteria were resuspended in 1 ml LB, diluted in 10-fold series from 10^-1^ to 10^-7^ and spread onto LB agar supplemented with chloramphenicol at 4 µg/l to select the first event of recombination. Transconjugants re-isolated on this medium were then spread onto LB agar without NaCl supplemented with 5% sucrose and incubated at 19°C for 2-3 days to allow the second event of recombination. Sucrose-resistant colonies were then patched on LB-Cm plate to check plasmid loss.

The in-frame deletion mutations ∆*ssmG* (NCBI Reference Sequence: WP_022634121.1;), ∆*oocL* (NCBI Reference Sequence: WP_023638021.1) and ∆*zsmA* (NCBI Reference Sequence: WP_022632849.1) were constructed in strain DS58 by using derivatives of the pRE112 suicide plasmids containing upstream and downstream 500-bp DNA of each gene (Table S3). In-frame deletions were checked by PCR as previously described. Then double mutants were constructed: DS58 ∆*ssmG* ∆*oocL* was obtained from the ∆*oocL* single mutant*,* DS58 ∆*ssmG* ∆*zmsA* from DS58 ∆*zmsA*, DS58 ∆*oocL* ∆*zmsA* from DS58 ∆*zmsA*. Then, a triple mutant DS58 ∆*ssmG* ∆*oocL ∆zmsA* was obtained from DS58 ∆*ssmG* ∆*zmsA*. The presence of double or triple deletions in the genome of the mutants was checked by colony PCR.

**Construction of ∆*ssmG*/*ssmG^+^,* ∆*oocL*/*oocL^+^* and ∆*zmsA/zmsA^+^* revertant strains.**

We first constructed complementation plasmids of the ∆*ssmG*, ∆*oocL*, and ∆*zmsA* mutants but they failed to restore the wild-type phenotype of two of the three mutants (data not shown). This could be explained by a polar effect of the mutations or the poor plasmid stability. Then, we decided to put back each wild type genes *ssmG^+^, oocL^+^ and zmsA^+^* in their original locus in the single deleted mutants. To do this, pRE112-*ssmG*^+^, pRE112-*oocL*^+^ and pRE112-*zmsA*^+^ were constructed by cloning after PCR and Gibson cloning each gene with their 0.5-kbp downstream and upstream DNA region into the pRE112 suicide plasmid (Table S1). The WT genes were put back to their chromosomal locus by homologous recombination following the standard procedures. Loss of the plasmid was checked and presence of the WT genes was verified by colony PCR by using the appropriate pair of primers (Table S3).

**Construction of the ∆*arcZ* mutants and the complementation plasmids.**

The ∆*arcZ* mutants of the WT *D. solani* strains D s0432-1 and IPO2222 were generated by allelic exchange using pRE112 as previously described, using oligonucleotide pairs L1333/L1334 and L1335/L1336 (Table S3). Since *arcZ* and *arcB* genes overlap, the last 14 bp of *arcZ* were conserved to avoid deletion of *arcB* 3’ end. Correct deletion was checked by colony PCR.

For the complementation assay, the low-copy plasmid pWSK29 was first made mobilizable by cloning a PCR-amplified oriT of SEVA plasmids [7] between the two SfoI restriction sites, leading to the pWSK29-oriT. The *arcZ*_Ds0432-1_ (*arcZ*1) or *arcZ*_G90A_ (*arcZ*2) alleles under the control of their own promoter were amplified by PCR with oligonucleotide pairs L1331/L1332, and then cloned into EcoRV-linearized pWSK29-oriT. Plasmids were checked by restriction map and DNA sequencing. Then, they were transferred to *D. solani* strains by mating.

**RNA isolation and Northern detection**

To assess ArcZ presence, bacteria were grown at 30°C in M63 medium with 1% sucrose to OD_600_ 1.5. Then 1 ml was fixed by mixing with 1 ml methanol at 4°C and centrifuged. Pellets were frozen at -80°C until RNA extraction. RNA extraction was performed by resuspending the bacterial pellets in 100 µl RNAsnap (18 mM EDTA pH 8.0, 0.025 % SDS, 95% formamide) and heated twice at 95°C for 7 minutes. The sample was then mix with 650 µl of Tri-reagent (Zymo Research) and RNA was extracted by following the protocol in the Direct-zolTM RNA MiniPrep kit (Zymo Research). Elution was made in 30 µL of Dnase/Rnase-free water. RNA concentration was determined by Nanodrop.

For Northern blot analyses, 0.8 µg of total RNA were loaded per lane of a 6% acrylamide-urea gel (Invitrogen), as well as 5 µl of Riboruler low range (Thermo Fisher Scientific) and 5 µl Low Range ssRNA Ladder (New England Biolabs) as ladders. After migration at 200V in TBE (Tris Borate EDTA) buffer, gels were soaked in TBE with ethidium bromide and imaged to check for RNA and migration quality. Gels were then transferred to a nylon membrane (Hybon-N+; GE Healthcare) by electrophoretic transfer in 0.5x TBE buffer (30 minutes at 300 mA). After crosslinking at 254 nm (Statagene, Stratalinker), the membranes were pre-hybridised for 4 h at 42°C in UltraHyb (Ambion) and then incubated at 42°C overnight with the biotin-labeled probe LA195 (anti ArcZ ; sequence AAAAAAAATGACCCCGACCGAGGTCGGGGTGCGCGAATTATG). Following washing at 65°C with SSC 2x SDS 0.1%, hybridized probes were revealed with the Chemiluminescent Nucleic Acid Detection Module (Pierce) which uses Streptavidin-conjugated HRP peroxidase and luminol. Luminescence signals were acquired using an imaging workstation equipped with a charge-coupled device camera (Thermo Scientific). ArcZ secondary structure was predicted using RNA-fold [8,9]

**Construction of the MLSA tree positioning strains within the *Dickeya* genus.**

Nucleotide sequences of 12 housekeeping genes (*fusA, gapA, glyA, groEL, gyrB, mdh, purA, recA, rplB, rpoD, rpoS*, and *secY*) were retrieved from 58 *Dickeya* genomes. Houkeeping genes were concatened, aligned, and the evolutionary history was inferred by using the Maximum Likelihood method based on the Tamura-Nei model [10]. The tree with the highest log likelihood (-81928,10) is shown. The percentage of trees in which the associated taxa clustered together is given next to the branches. Initial tree for the heuristic search was obtained automatically by applying Neighbor-Join and BioNJ algorithms to a matrix of pairwise distances estimated using the Maximum Composite Likelihood (MCL) approach, and then selecting the topology with superior log likelihood value. The tree is drawn to scale, with branch lengths measured in the number of substitutions per site. The analysis involved 58 concatened nucleotide sequences. Codon positions included were 1st+2nd+3rd+Noncoding. All positions containing gaps and missing data were eliminated. There were a total of 16596 positions in the final dataset. Bootstrap values were calculated from 1000 replicate iterations. Evolutionary analyses were conducted in MEGA7 [11].
