## Supplementary material for "A natural single nucleotide mutation in the small regulatory RNA ArcZ of *Dickeya solani* switches off the antimicrobial activities against yeast and bacteria": Table S3.docx

**Table S1: Bacterial strains and plasmids**

| Bacterial strain and plasmid | Description | Source |
| --- | --- | --- |
| **Strains** |  |  |
| *Escherichia coli* K12 |  |  |
| DH5α | *supE44 lacU169 (*Φ*80lacZ*∆ M15) *hsdR17 (rK mK ) recA1 endA1 gyrA96 thi-1 relA1* | Laboratory collection |
| DH5α λpir | λpir phage lysogen of DH5α | Laboratory collection |
| MFD*pir* | *RP4-2-Tc::(∆Mu1::aac(3)IV-∆aphA-·∆nic35-∆Mu2::zeo) ∆dapA::erm-pir) ∆recA* | [1] |
| MG1655 | *F^–^ λ^–^ ilvG^–^ rfb-50 rph-1* | Laboratory collection |
| *Dickeya solani* D s0432-1 |  |  |
| DS49 | *D. solani* D s0432-1 WT | Laboratory collection |
| DS50 | DS49 spontaneous Nal^R^ resistant clone | This study |
| DS58 | DS49 *glmS::Tn7-gent*, Nal^R^ Gm^R^ | This study |
| DS68 | DS58 ∆*ssmG* (∆BJD21_RS20030 of cluster A) | This study |
| DS69 | DS58 ∆*oocL* (∆BJD21_RS15005 of the oocydin cluster B) | This study |
| DS70 | DS58 ∆*zmsA* (∆BJD21_RS05130 of the zeamine cluster C) | This study |
| DS425 | DS58 ∆*ssmG* ∆*oocL* | This study |
| DS426 | DS58 ∆*ssmG* ∆*zmsA* | This study |
| DS427 | DS58 ∆*oocL* ∆*zmsA* | This study |
| DS352 | DS58 ∆*ssmG* ∆*oocL ∆zmsA* (named ∆3 mutant) | This study |
| DS428 | DS58 ∆*ssmG*/*ssmG^+^* revertant | This study |
| DS466 | DS58 ∆*oocL*/*oocL^+^* revertant | This study |
| DS429 | DS58 ∆*zmsA/zmsA^+^* revertant | This study |
| DS354 | DS49 *D. solani* D s0432-1 ∆*arcZ_1_* |  |
| *Dickeya solani* IPO2222 |  |  |
| DS45 | *D. solani* IPO2222 WT | [2] |
| DS353 | DS45 *D. solani* IPO2222 ∆*arcZ_2_* | This study |
| DS486 | *D. solani* IPO2222 WT | Strain LMG25993 of the BCCM collection |
| **Plasmids** |  |  |
| pRE112 | Suicide vector for allelic exchange, Cm^R^, *sacB*, *oriT* RP4, *ori*R6K | [3] |
| pTn7-M | Km^R^ Gm^R^, *ori R6K*, *Tn7L* and *Tn7R* extremities, standard multiple cloning site, *oriT* RP4 | [4] |
| pTNS3 | Ap^R^, *ori R6K*, *TnsABCD* operon, *oriT* RP4 | [5] |
| pEGL159 | pRE112-∆*ssmG* (∆BJD21_RS20030), Cm^R^ | This study |
| pEGL160 | pRE112-∆*oocL* (∆BJD21_ RS15005), Cm^R^ | This study |
| pEGL161 | pRE112-∆*zmsA* (∆BJD21_ RS05130), Cm^R^ | This study |
| pEGL325 | pRE112-*ssmG*^+^, Cm^R^ | This study |
| pEGL326 | pRE112-*oocL^+^*, Cm^R^ | This study |
| pEGL327 | pRE112-*zmsA^+^*, Cm^R^ | This study |
| pEGL302 | pRE112-∆*arcZ*, Cm^R^ | This study |
| pWSK29 | Amp^R^, pSC101 ori, lacZp expression vector | [6] |
| pEGL332 | pWSK29-oriT | This study |
| pEGL333 | pWSK29-oriT-ArcZ_2_ | This study |
| pEGL334 | pWSK29-oriT-ArcZ_1_ | This study |
