## Supplementary material for "A natural single nucleotide mutation in the small regulatory RNA ArcZ of *Dickeya solani* switches off the antimicrobial activities against yeast and bacteria": Table S4.docx

**Table S2: Oligonucleotides used in this study**

| Oligonucleotide | Sequence (5’-3’) | Use |
| --- | --- | --- |
| **L6** | gttttcccagtcacgac | To check cloning in the MCS of pEGL332 |
| **L7** | caggaaacagctatgacc |  |
| **L365** | cacagcataactggactgatttc | Primers to check integration of the miniTn7 into the attTn7 site of *D. solani*. |
| **L848** | atgtggcgctgatcaaaggc |  |
| **L762** | gttattggtgcccttaaacg | Primers to check the correct cloning into pRE112. |
| **L763** | gcatccaacgccattcatgg |  |
| **L828** | tgaactgcatgaattcccgggagagctcacgtaagcatcttcaggatgc | For amplification of the upstream and downstream 0.5-kb DNA fragments of *D. solani* *BJD21_RS20030* (*ssmG*) and their cloning into pRE112 |
| **L829** | gtccgtgcagacagatcgctcattctttatcctccg |  |
| **L830** | aagaatgagcgatctgtctgcacggaccattga |  |
| **L831** | ccgatcccaagcttcttctagaggtacccgtattgcgtcatcacctcac |  |
| **L832** | tgaactgcatgaattcccgggagagctcgccggattatccgttggatg | For amplification of the upstream and downstream 0.5-kb DNA fragments of *D. solani* *BJD21_RS15005 (oocL)* and their cloning into pRE112 |
| **L833** | cctgttccaacctggcctgtcatcgttgtctacc |  |
| **L834** | aacgatgacaggccaggttggaacaggagaaatccg |  |
| **L835** | ccgatcccaagcttcttctagaggtaccgcacgatcctcataggtgtcc |  |
| **L836** | tgaactgcatgaattcccgggagagctcatctggatgtggaaacctttgc | For amplification of the upstream and downstream 0.5-kb DNA fragments of *D. solani* *BJD21_RS05130* (*zmsA*) and their cloning into pRE112 |
| **L837** | aacggtcatttattccctaacatatattccctcttacacga |  |
| **L838** | atatatgttagggaataaatgaccgtttaattaaaacgagcc |  |
| **L839** | ccgatcccaagcttcttctagaggtacctggcgatatccgctatgactg |  |
| **L912** | acaggaagccggtgttaatg | Primers to check the correct deletion of BJD21_RS20030 (*ssmG*) in *D. solani* WT or the correct reinsertion of *ssmG* in ∆*ssmG* mutant |
| **L913** | atgtcatgccggttaatggt |  |
| **L910** | ggtacggattacggcttgaa | Primers to check the correct deletion of BJD21_RS15005 *(oocL)* in *D. solani* WT or the correct reinsertion of *oocL* in ∆*oocL* mutant |
| **L911** | gctgaataggacgcggataa |  |
| **L908** | cggtgggttttatttcatgg | Primers to check the correct deletion of BJD21_RS05130 (*zmsA*) in *D. solani* WT or the correct reinsertion of *zmsA* in *∆zmsA* mutant |
| **L909** | ccggtctttttcagaagtgc |  |
| **L1333** | aactgcatgaattcccgggagagctcggcgatacaaataaaccctattgg | For amplification of the upstream and downstream 0.5-kb DNA fragments of *D. solani* *arcZ* and their cloning into pRE112 |
| **L1334** | aaaaaaatgacccttacaagaatcattaaaatgtgtgatgtagattagt |  |
| **L1335** | tgattcttgtaagggtcattttttttcagcctctg |  |
| **L1336** | gatcccaagcttcttctagaggtaccacggcgaatgtgctcaaagat |  |
| **L937** | tgacccgcgaccagaccacgtttgcggccgcttttccgctgcataaccc | For amplification of the oriT region of pSEVA541 and cloning in pWSK29 |
| **L938** | gttgcagccctagatcggccacagcggccgtacggccagcctcgcagag |  |
| **L1331** | aggccgcctaggccgcggccgcgcgaattccactcaagcctcatttcattacatc | To amplify *arcZ* of *D. solani* with its own promoter and cloning in pEGL332 |
| **L1332** | gcggccgcaagcttgcatgcctgcagtgggtttcagaggctgaaaaaaaatgac |  |
